# Molecular architecture and dynamics of SARS-CoV-2 envelope by integrative modeling

**DOI:** 10.1101/2021.09.15.459697

**Authors:** Weria Pezeshkian, Fabian Grünewald, Oleksandr Narykov, Senbao Lu, Valeria Arkhipova, Alexey Solodovnikov, Tsjerk A Wassenaar, Siewert J. Marrink, Dmitry Korkin

## Abstract

Despite tremendous efforts by the research community during the COVID-19 pandemic, the exact structure of SARS-CoV-2 and related betacoronaviruses remains elusive. Being a key structural component of the SARS-CoV-2 virion, the envelope encapsulates viral RNA and is composed of three structural proteins, spike (S), membrane (M), and envelope (E), which interact with each other and with the lipids acquired from the host membranes. Here, we developed and applied an integrative multiscale computational approach to model the envelope structure of SARS-CoV-2 with near atomistic detail, focusing on studying the dynamic nature and molecular interactions of its most abundant, but largely understudied, M protein. The molecular dynamics simulations allowed us to test the envelope stability under different configurations and revealed that the M dimers agglomerated into large, filament-like, macromolecular assemblies with distinct molecular patterns formed by M’s transmembrane and intravirion (endo) domains. These results are in good agreement with current experimental data, demonstrating a generic and versatile integrative approach to model the structure of a virus *de novo*. We anticipate our work to provide insights into critical roles of structural proteins in the viral assembly and integration, proposing new targets for the antiviral therapies.

---

Betacoronaviruses are enveloped positive-strand RNA viruses. It has been nearly 20 years since the first outbreak of a pathogenic betacoronavirus, SARS-CoV, which was followed by MERS-CoV, and, most recently, SARS-CoV-2 (1-3). Yet, despite tremendous efforts in the elucidation of structural details of the viral proteins and protein complexes, a detailed virion structure of a betacoronavirus remains unsolved, due to the complexity and plasticity of these viruses. The virion particles of SARS-CoV-2 and closely related viruses are formed by four structural proteins: the spike (S), membrane (M), envelope (E), and nucleocapsid (N) proteins (4). The first three of these proteins oligomerize, and together with lipids from the host cell membranes, form the viral envelope, while the function of N protein is to organize, pack, and protect the viral RNA strand. Detailed structural knowledge of the viral envelope is critical because it allows for a mechanistic understanding of interactions between the virus and the host cell, and because the envelope surface presents potential drug targets for therapeutic interventions (5).

Electron microscopy (EM) and electron tomography (ET) studies of several betacoronaviruses, including SARS-CoV-2, have revealed that the morphology of the viral envelope is conserved while allowing certain flexibility in its overall shape (6, 7). These recent studies also suggest that the envelope forms an ellipsoid with the average diameters estimated for SARS-CoV-2 to range between 53-77 nm, 77-95 nm, and 85-109 nm, respectively; comparable sizes have been previously reported for SARS-CoV (8). The architecture of the envelope includes 26±15 S trimers (6), which is less than the previously reported number of trimers in a closely related mouse hepatitis virus, MHV (74 S trimers on average (8)). The number of M dimers in SARS-CoV-2 is currently unknown but is expected to be ∼1,100, comparable to the amount of M dimers for the envelopes of three other betacoronaviruses, SARS-CoV, MHV, and FCoV (8). Lastly, E pentamers are estimated to localize in small numbers in the envelopes of the above coronaviruses, including SARS-CoV-2 (9-11). Being the most abundant protein of the envelope, the M protein is integrated into the virus’ lipid bilayer in a homodimeric form and plays an important yet not fully understood structural role. For instance, it is still unclear whether M is sufficient to form a stable envelope structure in coronaviruses, or S and E are also required (12). The role and nature of the interactions between these three proteins were suggested to be more complex than originally expected (13). More recent work based on EM studies revealed the ability of M dimers to form lattice-like structures and highlighted the role of the endodomain of M in lattice formation and interactions with other structural proteins, including N (8, 14).

The recent EM and ET studies of the virion particles of SARS-CoV-2 (6, 7) together with earlier microscopy studies of SARS-CoV and MHV (8, 15) have provided important information on the morphology of the virion, including basic local and global geometric patterns formed by the structural proteins on the envelope surface, the stoichiometry of the structural proteins contributing to the envelope, and the distribution of S trimers. However, the details are obtained from averages of hundreds of images and thus provide more general, low-resolution, information on the spatial arrangement of the structural proteins. The main reason behind the lack of high-resolution imaging data for any coronavirus is the flexible nature of the viral envelope, which prevents fitting structures or structural models of proteins and lipids into the electron density. Thus, a *de novo* approach is required that does not rely on the density data (16).

An additional challenge for modeling the envelope is the lack of symmetry in the envelope, which is a typical feature of viral capsids. In addition, none of the three structural proteins constituting the viral envelope, E, M, or S, have been resolved experimentally in their full lengths for SARS-CoV-2. A full-length model of the spike trimer was recently constructed by combining high-resolution cryoelectron microscopy (Cryo-EM) data with the modeled structures of experimentally unsolved domains (17). A homology model of the E pentamer was also recently obtained (18), followed by refinement in a lipid bilayer (19), providing evidence for potential instability of the pentamer structure if structurally resolved as a single particle, without interaction with other structural proteins in the membrane (20). For the M dimer, however, no accurate model currently exists: the lack of previously detected homologous template structures prevents an accurate comparative modeling approach (21), while the top-scoring models of the two-domain monomer obtained with the state-of-the-art *de novo* protein structure prediction approaches (22, 23) could not be corroborated by the existing experimental evidence.

We developed an integrative approach to generate detailed models of the SARS-CoV-2 envelope by combining structural data from experiments and from homology models for the individual proteins, their oligomeric conformations, protein stoichiometries, the local geometries of the protein configurations, lipid bilayer composition, and the global geometry of the envelope (Fig. 1, Supp. Fig S1, Methods). The integrative computational modeling pipeline included five stages (Fig. 1): 1) structural modeling and refinement, including generation of a *de novo* structural model of the M dimer as well as a lipid fingerprinting analysis for all envelope proteins; 2) data integration, using a mesoscale simulation protocol to construct initial structural configurations of the envelope by taking as an input the refined structures of the three homooligomers, their stoichiometry, composition of the lipid bilayer, as well as the geometry and size of the envelope; 3) molecular assembly, converting the initial mesoscale models to a near-atomistic coarse-grain (CG) representation based on the Martini 3 force field (24) and testing for stability; 4) molecular dynamics (MD) simulation, generating long timescale dynamics of the stable envelopes; and 5) molecular trajectory analysis, involving the analysis of the obtained structural trajectories and comparison to experimental data.

**Figure 1.**
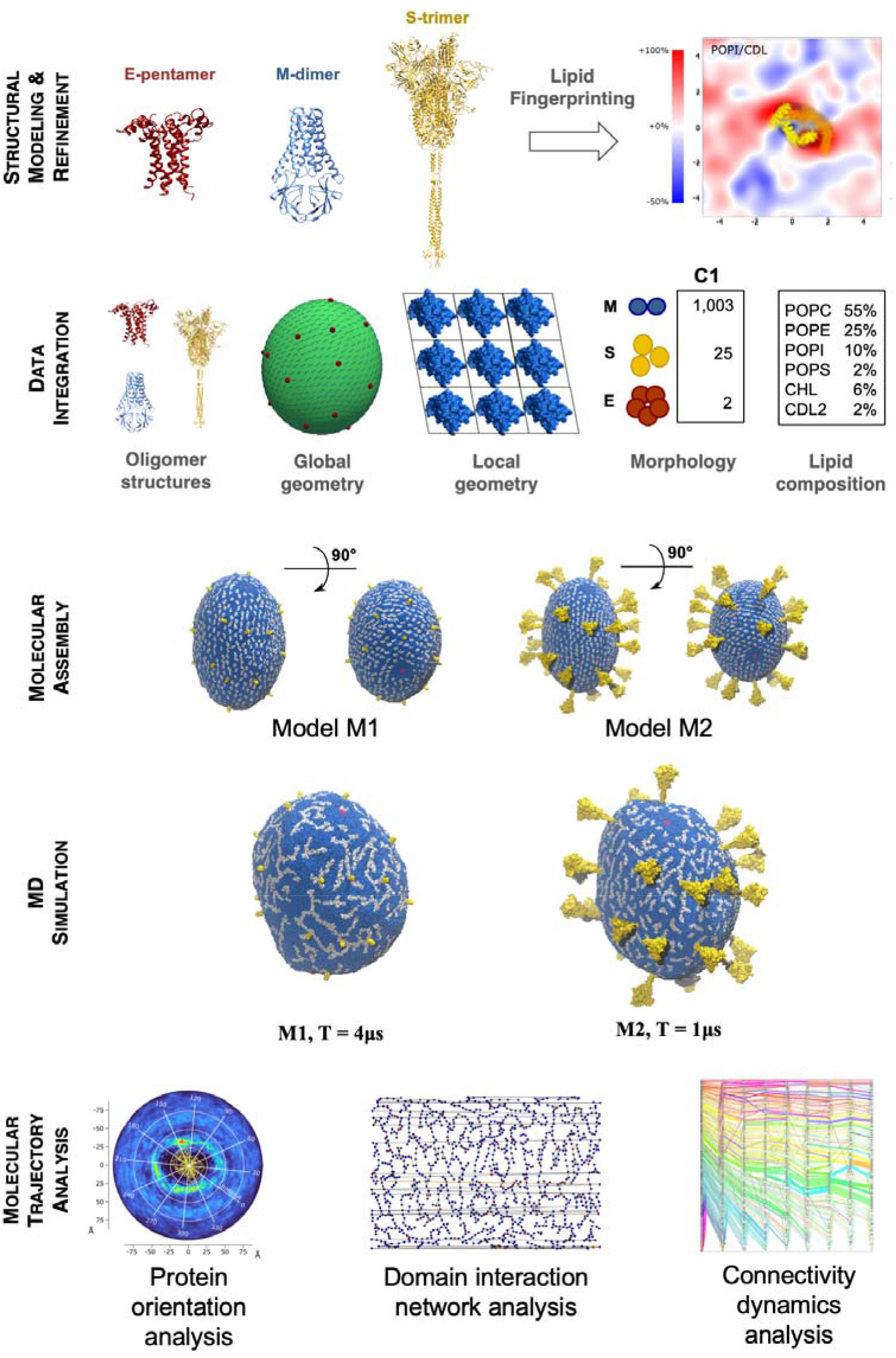
Key stages of integrative computational modeling pipeline: First, the structural characterization of S, M, and E proteins in both, monomeric and homooligomeric, forms was carried out, including a fingerprinting analysis to determine the enrichment of specific lipids around the proteins. Second, an initial structural configuration of the envelope was constructed by integrating data from multiple sources, captured in a mesoscale model and simulated with DTS. Third, multiple CG envelope models were assembled based on the configuration obtained from the mesoscale simulations, resulting in two stable models (M1, M2). Fourth, extended MD simulations of these stable models were performed. Last, structural and network analysis of the obtained structural trajectories was done to reveal characteristic patterns of the structural proteins in the envelope assembly.

Recently, a number of other computational models of the SARS-CoV-2 envelope have been developed. Voth and coworkers were the first to come up with a full envelope model, following a multiscale approach (25). The proteins and lipids were described at a supra-CG resolution, with their interactions calibrated with respect to all-atom reference simulations. A higher resolution model was developed by the Tieleman group (26), like ours, based on the Martini force field. However, a relatively small virus particle was simulated with a stoichiometry incompatible with current experimental data. Another Martini-based model was put forward by the group of Song (27). The model features N-bound RNA segments, and was back-mapped to full atomistic resolution. Unfortunately, only a short simulation was performed, preventing observation of structural changes. Our modeling approach is distinct in the sense that we use the most recent experimental data to build our envelope model, and we include an integrative modeling structure of the M-dimer that is more accurate compared to the *de novo* one. In addition, we probe the envelope dynamics over multiple microseconds allowing us to capture significant molecular rearrangements.

## Results and Discussion

### Modeling of M dimers

To enable CG simulations of the entire virion envelope, high resolution structures of the constituting S, E, and M proteins are required. The CG representations of the homooligomers of the structural proteins were obtained from available atomic models (S and E) and using a novel integrative modeling approach (M) (Fig. 2C, Supp. Fig. S2A,B). These were coarse-grained and then refined in the presence of a lipid bilayer using the Martini 3 force field (Methods). The initial model of the full-length S trimer was obtained previously using an integrative modeling approach (17), and a model of the E pentamer was obtained previously using homology modeling (18, 19). In contrast to the S and E proteins, a structure of M or its homodimer supported by experimental observations or evolutionary inference did not exist. Therefore, we first modeled the structure of M dimer using an integrative approach (Supp. Figs. S3-S6). The procedure started with the *de novo* modeling of the monomeric structure of M, followed by constraint symmetric docking to create a homodimer that satisfies the geometric constraints obtained based on 1) the envelope’s membrane thickness, 2) mutual orientation of the monomers, and 3) the approximate local geometric boundaries of a single M dimer complex, previously obtained from microscopy data of the SARS-CoV envelope (8). However, preliminary CG MD simulation of the envelope using the obtained top-scoring *de novo* model of the M dimer revealed the structural instability of the dimeric complex prompting us to the further refinement of the model.

**Figure 2.**
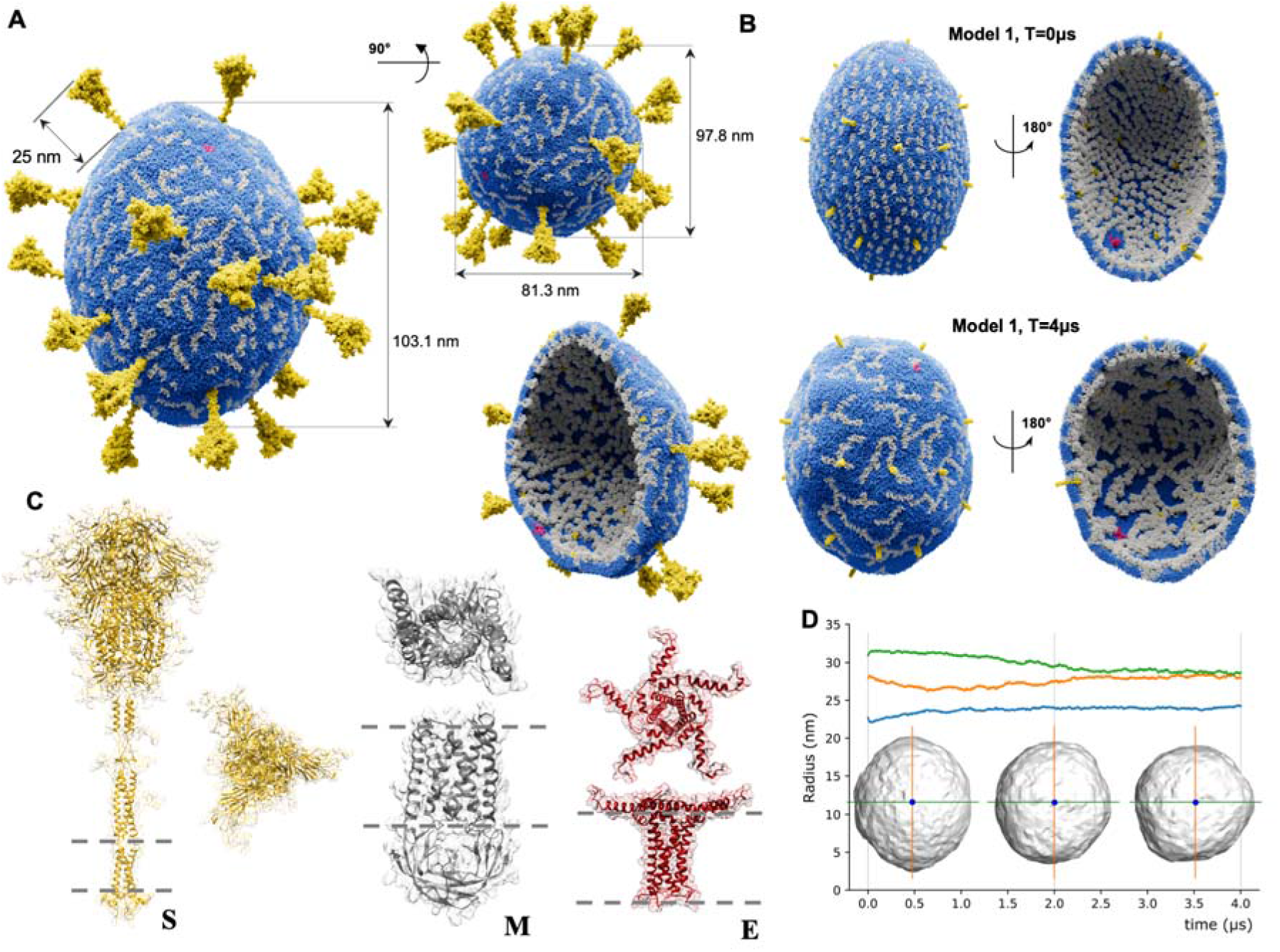
Structural characterization of SARS-CoV-2 viral envelope and its components: **A**. An envelope model (M2) obtained from molecular composition C1 (2 E pentamers, 25 S trimers, 1003 M dimers) and including full-length structures of S trimers after 1μs simulation run. Lipid molecules are depicted in sapphire blue, E pentamers in ruby red, M dimers in silver, and S trimers in gold. Principal diameters have values of 81.3 nm, 97.8 nm, and 103.1 nm. Height of the outer part of S protein is 25 nm. Surface of the envelope displays “filament” patterns formed by transmembrane domains of M dimers, while the internal part of the envelope shows tight packing of M dimers’ endodomains assemblies; **B**. Envelope model M1 from molecular composition C1 using truncated S trimer structures at the start of the simulations (top) and after 4μs (bottom); **C**. Structural proteins S, M, and E representing the main structural building blocks of SARS-CoV-2 envelop in their physiological oligomeric states, in side and top views: S trimer, M dimer, and E pentamer. The grey dashed lines correspond to the membrane boundaries. The structures are shown in different scales. **D**. Change of the viral shape during the simulation defined through the principal gyration radii. The two largest principal radii converge to the value ∼28 nm while the third one converges to ∼24 nm. The actual diameters of the model after 4μs simulation were 103.1 nm, 97.8 nm, and 81.3 nm, respectively.

We found that the *de novo* homodimer models of M that satisfy all the above constraints appeared to share striking structural similarity with another recently resolved homodimer of SARS-CoV-2, the ORF3a protein (28). The similarity included 1) the same two-domain fold composition and 2) the same combination of secondary structure elements as in our *de novo* model, but with a slightly different arrangement of the secondary structure elements in the transmembrane domain (Supp. Figs. S4, S5). We thus further refined the M dimer model by constructing a new structural template as a scaffold of the same secondary structure elements as in the original *de novo* model, each of which was structurally superposed against the ORF3a dimer. We then applied a novel integrative template-modeling protocol using the newly designed template and followed by a refinement protocol guided by the electron density of ORF3a (Methods, Supp. Fig. S6). The rationale for this approach was that the newly designed template, based on the ORF3A secondary structure topology and including the original secondary structure fragments of M dimer extracted from the top-scoring *de novo* model, would improve the arrangement of the secondary structure elements, making the model more stable, while maintaining the structural similarity with the original *de novo* model. The resulting model not only provided a tighter, more stable packing of M monomers in the dimer; but the shape complementarity of M dimers with each other allowed for a natural tiling of multiple dimers into the “filament” structures, consistent with the previously proposed model of M dimer lattices based on the microscopy study of SARS-CoV envelope (8, 29). Importantly, the *de novo* M dimer model proved stable in subsequent simulations of the full envelope. Furthermore, a 200 ns all-atom simulation of the two TM domains embedded in a lipid bilayer also resulted in a stable complex (Supp. Movie S1), while a 200ns simulation of the TM dimer of the top-scoring *de novo* model appeared to be unstable (Supp. Movie S2).

### Constructing the viral envelope

The modeling of the entire envelope started with the generation of a mesoscale model using dynamic triangulated surface (DTS) simulation (30, 31) on a triangulated mesh, matching the dimensions of the virion envelope. The mesh included a set of vectors each representing one protein and its orientation in the envelope surface. To set up the initial positions of the structural proteins in the envelope structures, available EM data was used only to obtain information on local geometry (Methods, Supp. Fig. S1). The global geometric patterns observed in the EM studies were not used during modeling, but only to evaluate our model (*vide infra*).

The DTS simulation provided us with an initial guess of the protein organization and orientation on the fixed geometry of the envelope. This model was subsequently backmapped using TS2CG to near-atomic resolution (32), based on the CG Martini 3 models of the proteins and lipids (24) with specified stoichiometries (Supp. Fig. S1). This resulted in an initial arrangement of the oligomeric protein structures embedded in a lipid bilayer comprising up to six types of lipid molecules (palmitoyl-oleoyl-phosphatidylcholine, POPC; PO-phosphatidylethanloamine POPE; PO-phosphatidylserine, POPS; PO-phosphoinositol, POPI; cardiolipin, CDL2; and cholesterol, CHL), with a composition reflecting that of the endoplasmic reticulum (ER), but also considering enrichment of specific lipids due to interactions with the proteins, based on the lipid fingerprinting analysis. Specifically, this analysis revealed an enrichment of anionic lipids around the proteins, slightly so for PS and most notably for the doubly charged cardiolipin (Methods, Suppl. Materials, Supp. Fig. S2C, Supp. Table S1, Suppl. Fig S7). This sequestering of negatively charged lipids is consistent with recent results of all-atom simulations in case of isolated M-dimers (33). To take the affinity for PS and cardiolipins into account, we added an increased percentage of these lipids in some of the final envelope models, leading to an overall lipid composition as specified in Table 1.

**Table 1.** Overview of system compositions. Shown are the compositional details of the two stable envelope models in molecular composition C1 (2 E pentamers, 25 S trimers, 1003 M dimers) with truncated (model M1) and full (model M2) S trimers. A CG water particle corresponds to 4 real water molecules.

| **Composition** | | C1, model M1<br>truncated S, 4μs | C1, model M2,<br>full S, 1μs |
| --- | --- | --- | --- |
| **Proteins** | S | 25 | 25 |
|  | E | 2 | 2 |
|  | M | 1,003 | 1,003 |
|  | **Total particles** | **968,373** | **1,237,398** |
| **Lipids** | POPC | 34,923 | 34,923 |
|  | POPE | 11,837 | 11,837 |
|  | POPI | 5,918 | 5,918 |
|  | POPS | 1,183 | 1,183 |
|  | CHOL | 2,663 | 2,663 |
|  | CDL2 | 2,663 | 2,663 |
|  | **Total particles** | **751,373** | **751,373** |
| **Solvent** | Na | 175,565 | 409,000 |
|  | Cl | 184,999 | 517,855 |
|  | Water | 13,045,399 | 34,236,835 |
|  | **Total particles** | **13,220,964** | **35,163,690** |

Several envelope models were thus built and subjected to MD simulations to test the stability of the envelope structure. The overall models prepared for the simulation consisted of 20-30 million CG particles, representing about 100 million heavy atoms. The models varied in several key parameters: (i) different protein-to-lipid ratios, (ii) different stoichiometries of the structural proteins, (iii) full or partial ectodomains of the spike trimer included in the envelope structure, and (iv) different lipid compositions. In total, three independent simulations turned out to be stable. For the unstable models, the integrity of the envelope surface became compromised (an example of unstable structure simulation is shown in Suppl. Movie S2). The selected stable models (M1-3), simulated for 1-4μs, included ∼1.0-1.2M protein particles, ∼0.5-0.8M lipid particles, and ∼13M-35M solvent particles (Fig. 2A,B, Table 1, Supp. Tables S2, S3, Suppl. Fig. S8, Suppl. Movies S3-S5). A 4μs simulation took ∼1,560,000 CPU hours to compute on the TACC Frontera supercomputer.

We found that each of the key parameters played a role in the simulation. First, when selecting between two different protein-to-lipid particle ratios, the higher ratio value of 2.36 resulted in unstable structures, while the ratio of 1.44 resulted in a structural model that remained stable (models M1, M2). Given that the molecular composition was the same in both models (1,003 M dimers, 25 S trimers, and 2 E pentamers; we refer to this molecular composition as C1), the different ratios were due to the different lipid numbers (36,645 and 60,141 molecules, respectively), suggesting that the lipid concentration plays a role in the envelope stability, a finding supported by recent CG simulations of cell-scale envelopes (34). Another factor that affected the model stability was a higher number of solvent particles, compared to the stable models, leading to pore formation and subsequent membrane rupture.

Second, we found that varying the stoichiometries of E pentamer, S trimer, and M dimer under the same conditions does not affect the envelope stability. For instance, when we significantly increased the proportion of S trimers (truncated form) creating a model in an oligomeric composition C2 that included 3E pentamers, 71 S trimer, and 1080 M dimers (Supp. Table S3), while maintaining the same protein-to-lipid ratio and the number of solvent particles as in models M1 and M2, we found that the new model, M3, was also stable after 4μs simulation (Supp. Fig. S8). The stability of viral envelope with different stoichiometries is in line with the experimental evidence suggesting a range of different stoichiometries to be found *in vivo*. The behavior of the envelope model M2 that included the full-length S trimers and composition C1 was similar to the ones of the truncated model M1 (Fig. 2A,B, Supp. Fig. S2B): a 1μs trajectory of the former was comparable to the first 1μs of the 4μs trajectory for the latter. Finally, variations in lipid composition did not appear to impact the stability of the envelope.

### Plasticity of viral shape

In the 4μs simulations, we consistently observed changes in the viral shape (Fig. 2D, Supp. Figs. S9, S10), with the initial diameters of the ellipsoid of a model in composition C1 changing from *d*_1_ ≈ 109.9nm, *d*_2_ ≈ 97.8nm, and *d*_3_ ≈ 76.2 at t = 0μs to *d*_1_ ≈ 103.1 nm, *d*_2_ ≈ 97.8 nm, and *d*_3_ ≈81.3 nm at the end of the simulation, t = 4μs. The obtained diameters were close to the range observed for the particles from the Cryo-ET images of SARS-CoV-2 (35). We also note, that while the experimentally observed structures of the virions had variable shapes, from a nearly spherical shape to a significantly elongated ellipsoid, the average shape of SARS-CoV-2 according to the Cryo-ET study is an elongated ellipsoid, not a sphere, hence the rationale for our initial model dimensions. The elongated shape of a virion particle was also reported in previous studies of SARS-CoV and the related betacoronaviruses (8). The calculated *d*_MAX_/*d*_MIN_ ratio of 1.27 for our final model falls within the range of average ratios observed in CryoET of the SARS-CoV-2 virion (35). Interestingly, the changes in the shape did not have significant effects on the surface area of the envelope (1.3% reduction) or its volume (0.6%) (Supp. Figs. S9, S10). Along with the diameters of the envelope shape, the principal radii of gyration were also converging, reflecting shape stabilization (Fig. 2D). Furthermore, analysis of the temporal changes of the viral dimensions together with the connectivity patterns of the envelope proteins suggested the presence of two distinct concurrent relaxation processes, separately affecting the two smallest and the two largest diameters. Specifically, we observed a faster process (0−1μs), followed by a slower process (0.5−4μs). During the faster process, the minor circumference (principal radii *r*_2_ and *r*_3_ corresponding to diameters *d*_2_ and *d*_3_) became more circular while during the slower process, the major circumference (principal radii *r*_1_ and *r*_2_ corresponding to diameters *d*_1_ and *d*_2_) also became more circular, thus making the minor circumference to become more elliptical again (Fig. 2D).

### Formation of patterns of M dimers

Our envelope models and the time scales of the simulations allowed a detailed assessment of the interactions between the different constituents, in particular those involving M dimers. To characterize these interactions, we focused on the preferential relative orientations of protein neighbors and second neighbors (Fig. 3A-C), which were determined using a method for orientation analysis (36). The results showed that the M dimer transmembrane domains (TMDs) preferentially formed filament-like assemblies, without contacts between adjacent filaments (Figs. 3A,B). In contrast, the endodomains (EDs) appear tightly packed, binding neighbors in two directions (Figs. 3A,B), thus showing a tendency for the formation of a well-ordered lattice. Intriguingly, the lattice vectors of the TMD domains extracted from M dimers in our model were identical to the ones extracted from the previously reported EM data (8). Combining the (averaged) relative orientations of the TMDs and the EDs with the projected densities of the proteins revealed the characteristic patterns of densities. Specifically, the averaged orientations of TMDs fit almost exactly with the ‘lattice’ patterns previously observed from the averaged Cryo-EM virion structures of SARS-CoV and related betacoronaviruses (Fig. 3A,B) (8), despite the fact that this information was not used during construction of the models. Even the characteristic lack of density in the unit cells’ corners previously observed in MHV and SARS-CoV was clearly noticeable in our data (Fig. 3B). These patterns were consistent in the models with both compositions, C1 and C2. The orientations of EDs revealed a different kind of pattern, compared to the one of TMs. Specifically, our model showed that the ED dimers formed triangulated structures, a pattern common in engineering rigid frame structures (37). Unlike TMs, the formation of EDs have not been experimentally characterized by microscopy before, because it can only be observed from the virion’s interior. To ensure that the observed property of M dimers forming the filament-like assemblies was not a consequence of the initial setup, we have performed additional simulations of a subset of randomly placed and oriented 41 M dimers in a flat bilayer system with the same lipid composition as the full envelope simulations (Methods). After a 13 μs simulation, we observed that the M dimers formed filaments that were reminiscent of the ones we observed in the full envelope simulations (Suppl. Fig. S11, Suppl. Movie S6). In contrast to the strong preferential orientation of M dimers, the orientation of M dimers around S trimers did not show clear preference in attachment (Fig. 3C), which could be due to the stronger orienting effects of M dimer interactions as well as to the worse statistics for the interactions between S and M.

**Figure 3.**
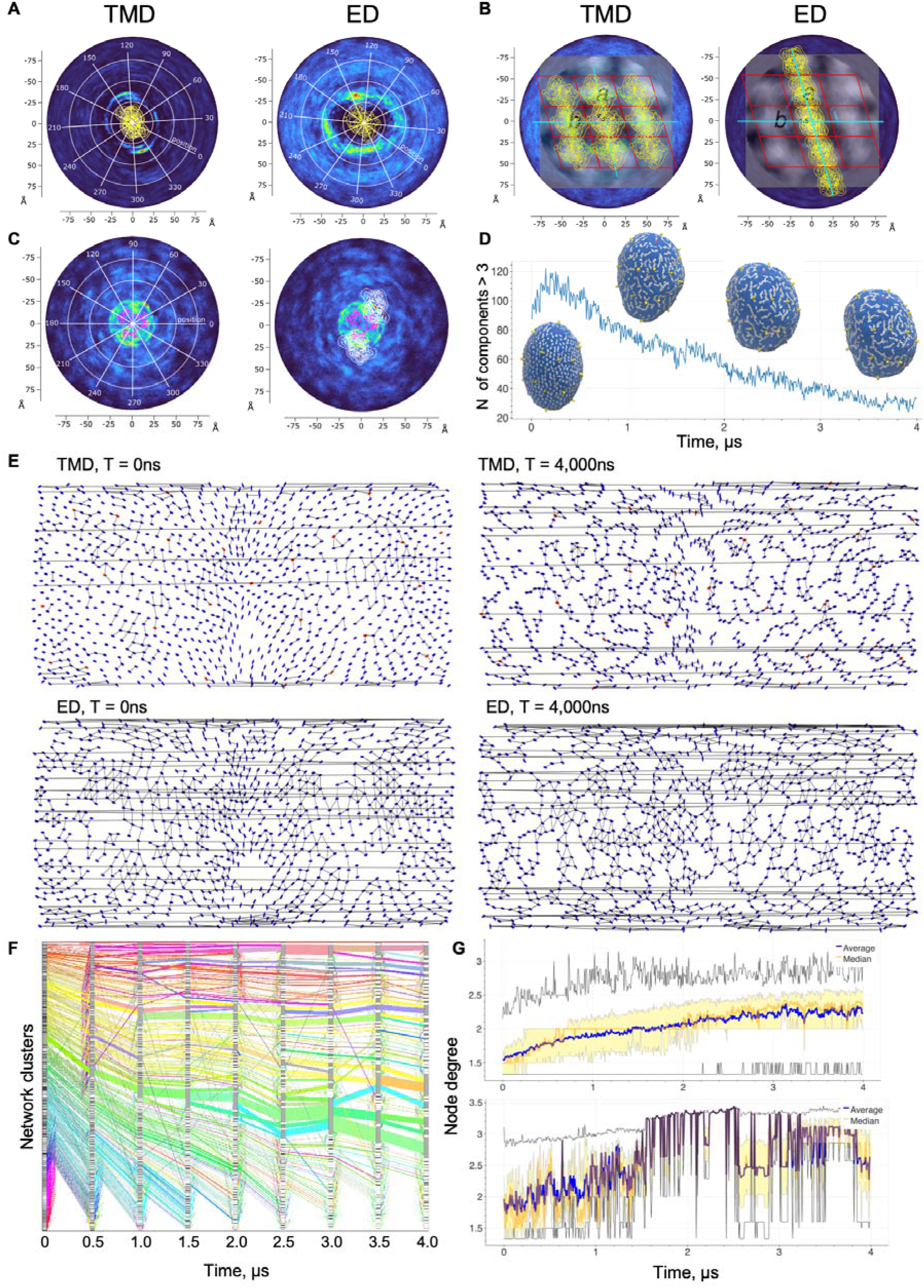
Structural and network analysis of the envelope assembly. **A**. Orientation preference of the transmembrane domains (TMD) and endodomains (ED), two main components of the M proteins in their dimeric state. TMD components display two preferred locations at 135° and 315°, ED components display four preferable locations at 135°, 315°, 225° and 75°; **B**. Super position of averaged Cryo-EM images previously obtained from SARS-CoV envelope with M dimer models obtained separately for TMDs and EDs and arranged according to the preferred interaction positions from panel A, demonstrating a near-perfect correspondence between our model and the Cryo-EM images. **C**. Orientation analysis of the contacts between S trimers and M dimers. Shown are positions of the S trimer (pink), the density of positions of M dimers (left panel) and the same density together with two M dimers (white) positioned around two distinct high-density locations at 60° and 260° (right panel). **D**. The number of connected components that have at least three nodes dynamically change during the simulation; included into the same plot are envelope models at t=0μs, 1.5μs, 2.5μs, and 4μs in molecular composition C1. **E**. Mercator projection of physical domain-domain interaction network established for molecular interactions of M dimers (blue ellipses) with each other and with S trimer (red ellipses) for the models in molecular composition C1. M dimer’s TMDs and EDs are arranged into separate domain-domain interaction networks. The orientation of ellipses corresponds to the orientation of the corresponding oligomers. An ellipse with the major axis positioned horizontally corresponding to the canonical orientations of the oligomers, as defined in panels A and C, for M dimer and S trimer, respectively. Connectivity increases after 4μs of simulations for both TMD and ED components; **F**. Sankey diagram showing clustering dynamics of the TMD components over the time for the model in molecular composition C1. Each of the displayed “flows” contains at least 25 network nodes. One can see a drastic increase in the cluster size for a small number of connected components as the simulation progresses; **G**. Average node degree dynamics during the simulation for TMDs (top) and EDs (bottom). Yellow bands indicate values for the 2^nd^ and 3^rd^ quantiles, black lines denote the minimum and maximum values. There is a clear trend for the increase of the node degree value for TMD components; the value plateaus during the last microsecond (3-4μs) around the value of 2.3. EDs’ node degrees have a much smaller spread and tend to converge to the value of 3.2.

To further characterize formation of the higher-order assemblies in the envelope through interactions between M dimers, a temporal analysis was performed of the domain-level physical interaction networks between the TMDs and EDs of M dimers and the transmembrane domains of S trimers. This analysis further illustrated the strikingly different nature of the M dimer’s key domains (Fig. 3D-G, Supp. Fig S12-S15), supporting our previous findings. Throughout the simulations, the domain interaction networks appeared to undergo drastic rearrangement via two distinct phases, with the number of connected components of three and more dimers first rapidly rising during 0-200ns up to ∼160 components and then slowly saturating to ∼30 components (Fig. 3D). The total number of connected components closely follows a bi-exponential law (Suppl. Figs. S12, S13), suggesting two processes running concurrently: a faster local rearrangement and a slower filament assembly. The two-process formation of the connected components was also evident from the clustering analysis (Fig. 3F), indicating the initial formation of many small clusters, followed by the preferential growth of the largest connected components. It was also interesting to see that the second, slower process of growing connected components started before the first, faster stage of forming initial small assemblies of M dimers was over. The analysis of the interaction network of TMDs also supported the formation of filaments observed on the surface of the envelope (Fig. 3E); the network revealed that these filaments were occasionally connected even further into larger assemblies. In contrast to the filament-like network topology, the EDs network consisted of connected triangulated components, which may contribute to the structural rigidity of the envelope. The difference between the TMD and ED network topologies also followed from the temporal node degree distributions for each network: the average node degrees converge to ∼2 in TMD network and ∼3 in ED network (Fig. 3G). Lastly, all other major network parameters appeared to converge to the stable values (Suppl. Figs. S12-S15), thus further drastic changes in network topologies were not expected.

## METHODS

### 1. Modeling and structural refinement of M dimer

Unlike for proteins S and E, no experimental structures have been solved for SARS-CoV-2 M protein or any of its homologs, neither as a monomer nor as a dimer, which is its physiological conformation in the envelope. Thus, a comparative modeling approach cannot be applied, and a novel integrative approach was introduced that utilized geometric constraints of the M dimer in the envelope derived from the low-resolution CryoEM images of the envelopes of the closely related coronaviruses, SARS-CoV and MHV, as well as a high-resolution CryoEM structure of a dimer for another SARS-CoV-2 protein sharing substantial structural similarity (see Supp. Figs. S3, S4).

The approach included six steps. First, an ensemble of models of M monomer was obtained using *de novo* modeling methods AlphaFold and I-TASSER (22, 38, 39). The top 5 models from each method were then selected, and 200 models of M homodimers overall were obtained using two symmetric docking approaches, SymDock and Galaxy (40-42) (10 docking models for each of the 10 monomers for each docking approach). Next, a set of geometric constraints was applied to an ensemble of the 100 top-scoring homodimer models (50 for each symmetric docking). The geometric constrains include (1) the dimer axial dimensions and the shape of the part of the packaged dimer located on the envelope surface, (2) the orientation of the monomers in the membrane, and (3) the approximate dimensions of the transmembrane domain (TMD) of a single packaged M dimer defined by the envelope membrane’s thickness. Specifically, from the previous analysis of SARS-CoV envelope (8, 15), it follows that TM domains of an average M dimer form a parallelogram, with rough dimensions between the two centers of adjacent dimer parallelograms measured to be 6.0 nm and 7.5 nm (Fig. 1). As a result, we filtered out those M dimer models whose TM domains would not fit into a parallelogram-shaped grid of these dimensions. We note that we did not require the TM domains to form a parallelogram-like shape, rather these constraints primarily affected the length of the modeled helices, filtering out models with significantly elongated one or two helices. Furthermore, the dimers were required to have N-termini of both monomers located on the exterior surface of the virion’s envelope, and the thickness of the envelope’s membrane was set to be equal to 4 nm (35, 43).

The analysis of the three best-scoring M dimers that satisfy all above geometric constraints provided us with an interesting finding: all three dimer models share some striking structural similarity with the ORF3 protein of SARS-CoV-2 (Supp. Figs. S4, S5), whose CryoEM structure was recently solved (28). In particular, we found that the *de novo* modeled structure of the endodomain of M and the experimentally obtained structure of C-terminal domain of ORF3 were structurally similar, while the transmembrane domain structures of M and N-terminal domain of ORF3 shared the same secondary structure elements (three helices) of the same lengths while the mutual arrangement of the helices was somewhat different, when comparing monomeric or dimeric structures (Fig. S16). Specifically, in all three initial models of M monomer, the second and third helices of the TM domain were aligned to the corresponding helices in one chain of ORF3a, while the first helix of the M monomer was aligned with the first helix of the other chain of ORF3a, suggesting a swapped arrangement in a dimer. This difference of the secondary structure arrangement in the TM fold decreases the dimerization interface surfaces in all three *de novo* generated models compared to ORF3A, which could account for the lack of stability observed in the models. Therefore, we hypothesized that M dimers and ORF3a were structural homologs, and the model of the M dimer could be further refined using the structural information from ORF3a dimeric structure (Fig. 1B).

To use the structural information from ORF3a, we first created a “fragmented scaffold’ structural template by individual structural alignment of the helices and the endodomain of the M dimer with the corresponding helices and endodomain in the ORF3a template structure (PDB ID: 6XDC). Then, we obtained a preliminary comparative model using the newly created fragmented scaffold template of M dimer. We note that, unlike a traditional template-based approach, here our template includes only a geometric arrangement of the six helices (three for each monomer structure) in a dimer. The structurally unstable, loop, regions of ORF3A were not used in the template and instead were modeled *ab initio* as a part of model optimization in MODELLER (44). The obtained M dimer model was refined using a protocol similar to the one used to obtain a full-length model S trimer (Fig. S5) (17). First, the linker regions of the two TMDs in the obtained comparative model of M dimer were refined by energy minimization, followed by refinement of the whole TMDs using Molecular Modeling Toolkit (45). Next, the overall M dimer structure was minimized using the CHARMM36 force field in GROMACS (46). We then placed the M dimer model into the experimental EM density map of the ORF3a dimer (EMD-22139 (28)) using Phenix (47) and relied on the EM map to further refine the structure in ISOLDE (48), a package for UCSF Chimera X molecular visualization program (49). ISOLDE uses OpenMM-based interactive molecular dynamics flexible fitting (50) using AMBER force field (51) and allows for the real-time assessment and validation of the geometric clashing problems. Each residue of the M dimer model (1-946) was then inspected and remodeled to maximize its fit into the density map. We considered both a deposited electron-microscopy map and a smoothed version with a B-factor of 100 Å^2^ as proposed in (17), but in our case there was no significant difference between these two versions of the refinement protocol.

To further compare the top-scoring *de novo* M dimer model and the one generated using our integrative template-based protocol, we studied the trajectories obtained via all-atom energy minimization. The overall M-dimer structure was minimized using the CHARMM36m force field in GROMACS (46). Then the structure was put into a realistic lipid bilayer using CHARMM-GUI Membrane Builder tool (52). The systems were neutralized with the addition of counter ions, then 0.15 M NaCl was added. Each simulation was run for 1mus. Principal component analysis (PCA) was performed on the resulting protein trajectories using an in-house developed plugin for Pymol to investigate and compare the dynamics of the different models and assess the stability.

### 2. Overall stoichiometry of the envelope

The molecular composition of the SARS-CoV-2 envelope and the stoichiometries of the structural proteins have been determined based on the current information obtained from CryoEM experiments, by analyzing the stoichiometries of other betacoronaviruses, and through arranging the three structural proteins into a geometrically constrained shape of the envelope. Comparable numbers of S trimers have been previously reported across other beta coronaviruses. An early model of BECV reported up to 110 S trimers (53), while another early model of TGEV provided a rough estimate of 100-200 S trimers (10). SARS-CoV has been determined to contain ∼90 copies of S trimers (8) per particle. Interestingly, the number of S trimers in a SARS-CoV-2 recently obtained from the cryoelectron tomography experiments was substantially less, namely 26 ± 15 S trimers (6). A similar estimate of 24 ± 9 S trimers was reported independently (54). The numbers of M molecules per virion are also consistent across studied betacoronaviruses. For SARS-CoV, MHV and FCoV, it was estimated at 1,100 M dimers on average per particle (8). Finally, E pentamers has been reported to localize in the envelope membrane in minute amounts (8, 9). In TGEV, 15-30 molecules of E protein, which corresponds to 3-6 pentamers was predicted (10). Similar numbers of ∼20 copies of E molecules, which correspond to four pentamers have been reported for MHV (11).

In our two main models, M1 and M2, we considered a molecular composition C1 that included 1,003 M dimers (2,006 monomers), 25 S trimers (75 monomers), and 2 E pentamers (10 monomers). M1 had truncated S trimers (for more details see the next section), while M2 had the full-length spikes. The number of E pentamers was slightly reduced from the estimates for other coronaviruses in order to maintain a E:S molar ratio close to the one reported for TGEV (10). In addition, to explore if the differences between the structural proteins of SARS-CoV and SARS-CoV-2 do not allow to maintain an envelope structure with a higher number of spike proteins for the SARS-CoV-2 virion, we have also simulated another model, M3, with a composition, C2, more similar to that one estimated or SARS-CoV (8) than to SARS-CoV-2 (54) that included 1,080 M dimers, 71 S trimers, and 3 E pentamers.

### 3. Martini3 protein models

The coarse-grained protein structures and input parameters were created from the atomistic reference structures of oligomers using martinize2 (55), following Martini3 guidelines for creating protein input parameters (56). The parameters for M protein were based upon our own refined atomistic structure, whereas parameters for E and S proteins were derived from the previously published models (17, 19). An elastic network was used to keep the secondary structure of the proteins fixed as required in CG simulations with Martini3. For the E protein, we created a custom elastic network, where the transmembrane domains of all 5 monomers were connected to keep the channel’s stability. In addition, the elastic network of the intracellular domains was applied only within each monomer to allow for flexibility. For the S protein, two CG models were created: (1) a model that included copies of the whole S protein together with glycosylation, and (2) a model that included a truncated structure of S consisting of HR2, TM, and CT domains in close proximity to the envelope (resid 1172-1273). Coordinates and parameters for the glycosylation of the full S protein were generated in a separate step using the polyply program (57) and a Martini3 beta sugar model with adopted parameters for the final Martini3 version. Composition of the glycans and attachment sites were considered the same as in the previously published work (19), except for the O-glycan, which was omitted due to missing parameters. No elastic network was applied to the glycans. Note that the previous tendency of Martini membrane proteins to aggregate too strongly has been remedied in version 3, as shown for a number of example cases in Souza *et al*. (24), and also in a dedicated independent test study (58). The aggregation of M dimers observed in our SARS-Cov2 envelope models is thus not an artefact of the chosen force field, but arises as a natural consequence of the interactions between the proteins.

### 4. Lipid fingerprinting analysis

While the lipid composition of the SARS-CoV-2 envelope has not been determined, it has been approximated as being close to the composition of the endoplasmic reticulum (51, 59). Membrane proteins are known to collect certain lipids in their immediate lipid shell. To analyze if any of the structural proteins had specific lipid interactions or favored lipid environments, the so-called lipid fingerprint analysis was conducted (60, 61). Specifically, we determined the lipid fingerprint under the consideration of clustering of M dimers in a model lipid membrane mimicking the composition of the endoplasmic reticulum (ER). Due to the fact that M dimers were found to also cluster in the flat membranes (Supp. Fig. S11), the lipid fingerprint was extracted from simulations with four M dimers after they formed clusters in the model membrane. For lipid fingerprint analysis of E pentamer and S trimer, each of the above homooligomers was added to the previous system of four M dimers (Supp. Table S1). Lastly, a simulation with a single M dimer was run to see if that has any effect on the results.

The additional factor that we investigated was the influence of cardiolipin on the lipid fingerprint. While a model of ER composition contains cardiolipin, the charge state of cardiolipin has not been specified. Here, we used the doubly charged version cardiolipin to see if this would have a strong effect on the lipid fingerprint, and we also ran a set of simulations without cardiolipin. For all systems, the lipid fingerprint analysis was analyzed per protein-multimer, but averaged over the copies in the cluster, accounting for the clustering properties of the M dimer. The composition of the bilayer was POPC (0.55), POPE (0.25), POPI (0.10), POPS (0.02), cholesterol (0.06), and cardiolipin 2 (0.02), which approximated the composition found in the ER (59). To account for the specific lipid binding of the E pentamer, it was placed with its immediate lipid shell as obtained from a separate free simulation. Each of the systems was run for 4μs, following the default simulation setup for Martini membranes (62). Then, the depletion enrichment index and a 2D density map was computed by analyzing the lipid compositions as a function of distance to the protein over time.

Based on the results, we slightly adjusted the lipid composition of the viral envelope following the reasoning that the high concentration of M dimers likely leads to an enrichment of lipids preferentially found in the M dimer lipid shell.

### 5. Building an initial CG structure of the full envelope

To build the initial CG structure of the envelope, we used a top-down protocol. First, by performing two types of Monte Carlo moves (vertex move and link flip (30)) on a spherical triangulated surface (TS) containing 1,030 and 1,154 vertices for models M1/M2 and M3, respectively, we drove the system to reshape to an ellipsoidal TS with aspect ratio of 0.67:0.89:1 to match an average shape reported in experimental measurements (54). Next, we assigned the location of each vertex of the TS to a homo-oligomeric complex (M dimer, S trimer, or E pentamer). The vertices are assigned so that the spatial distribution of complexes is uniform. To find a proper orientation of the oligomeric complexes, we ran a Monte Carlo simulation on the orientation of these complexes using a dynamic triangulated surface (DTS) simulation method (30) without updating the TS shape. To do so, we assumed that each homo-oligomer interacted only with the neighboring protein complexes by a potential defined as:

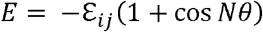

where *ε*_*ij*_ is the interaction strength, *N* is the least common multiple of the degree of the *i,j*-th complex symmetry in the plane of the membrane and θ is the angle between the inclusion vector (a vector representing the in-plane orientation of a homo-oligomer) residing on vertex *i* and the inclusion vector residing on vertex *j* after parallel transport to vertex *i* (63).

The final DTS output was then converted to a Martini model using TS2CG software (32). The rescaling factor in TS2CG converts the DTS unit length to nm (see (32) for more details). We choose this factor as small as possible to maximize the protein to lipid ratio of the envelope while generating a numerically stable system. Interestingly, this also gave us an envelope size within the range of experimentally reported dimensions. We created envelopes with different sizes (85,90,95,100,105,110 nm for the longest principal axes) and performed 100 steps of energy minimization using soft-core potentials (64), followed by a standard energy minimization for 10k steps and 1k steps molecular dynamics run (while proteins backbone and lipids headgroup were position restrained) to relax the lipid chains. We performed these steps without solvent particles. Only the system with the size of 110 nm went through these three steps; the rest were crashing due to high potential energies of bad contacts.

Next, the stable envelope particle was solvated by propagating an equilibrated Martini water box in the system and removing any water particle within a certain cutoff from the envelope particles. We aimed at finding the smallest cutoff length to avoid particle deformation due to an imbalance of surface area to volume ratio. It may appear as the cutoff could be just the Martini bead size. But this assumption does not hold for this system. The reference value of the area per lipid (APL) that we used in TS2CG was obtained from a smaller, flat system in which the protein concentrations were lower than in case of the envelope. These APL values can be different in the envelope due to the high curvature and high concentration of the proteins. Even a tiny change in APL could have a significant impact in the overall membrane area of this large system. Also, note that the volume scales as R^3^ while the area scales as R^2^. Additionally, TS2CG considers a uniform thickness for the envelope, while this might not be true for lipids close to the proteins. The minimum cutoff was obtained by finding stable envelopes (see an example of the simulation of an unstable structure in Suppl. Movie S2) after 300 ns simulations. We solvated the system and added Na+ and Cl-to neutralize the system, with an overall ion concentration of 0.9%. Finally, another energy minimization round is implemented followed by a short equilibration MD simulation run in the NVT ensemble and a longer run in the NPT ensemble using a Berendsen barostat. The resulting arrangement is approaching a limit of the physically possible fit in terms of the protein density with respect to the virion surface. The final production run was performed in the NPT ensemble, using the Parrinello–Rahman barostat with reference pressure of 1 bar and compressibility of 3×10^−4^ bar^−1^, respectively. The systems were equilibrated first in an NVT, followed by NPT ensembles using the Berendsen barostat and the temperature was kept constant at 310 K.

All CG-MD simulations were performed using the GROMACS package (version 2019) and the Martini 3 force field (24, 65). A time step of 20 fs was used, though for certain steps it was necessary to perform runs with shorter time steps (see 9-step protocol below). Both van der Waals (cutoff) and Coulomb interactions (reaction-field) were set to zero at 1.1 nm using the Verlet cutoff scheme, following recommended values for Martini-based simulations (66). Coulomb interactions were screened by a relative permittivity constant of 15.

The lipid composition of the viral envelope was taken to be the same as the one used in the lipid fingerprint simulations, since no significant lipid preferences were found for the M protein during the lipid fingerprint analysis, apart from a slight enrichment of the anionic lipids. Likewise, no significant enrichment around the S trimer was observed (Supp. Table S1, Supp. Fig. S7). To account for the specific lipid binding around the two E pentamers, each E pentamer was placed on the envelope together with its immediate lipid shell. This resulted in the final lipid bilayer composition of 39,755 POPC, 13,191 POPE, and 7,195 POPI lipids for the M3 model and an expanded set of 34,923 POPC, 11,837 POPE, 5,918 POPI, 1,183 POPS, 2,663 CHOL, 2,663 CDL2 for the final models of M1/M2.

### 6. Nine-step protocol with parameter settings

1. Energy minimization with softcore potentials: 100 steps, 0.02ps time step, flexible water, steep integrator, V-rescale temperature coupling, 310K temperature, no pressure coupling.
2. Energy minimization with positional restraints: 10,000 steps, 0.02ps time step, flexible water, steep integrator, no temperature and pressure coupling.
3. Equilibration with positional restraints: 10,000 steps, 0.001ps time step, rigid water, sd integrator, V-rescale temperature coupling, 310K temperature, no pressure coupling.
4. Solvation with 15 angstrom distance to the protein and lipids, and 0.9% salt content.
5. Equilibration with positional restraints: 100,000 steps, other parameters are identical to 2)
6. NVT equilibration with positional restraints: 500,000 steps, 0.002ps time step, rigid water, sd integrator, rigid water, V-rescale temperature coupling, 310K temperature, no pressure coupling.
7. Short run with positional restraints: 750,000 steps, 0.02ps time step, rigid water, md integrator, V-rescale temperature coupling, 310K temperature, Berendsen barostat, isotropic pressure.
8. NPT equilibration (no restraints): 100,000 steps, 0.01ps time step, rigid water, sd integrator, V-rescale temperature coupling, 310K temperature, Berendsen barostat, isotropic pressure.
9. Production run: 50,000,000 steps, 0.02ps time step, md integrator, Berendsen/Parinello-Rahman barostat (first and second run correspondingly), isotropic pressure.

Production runs for the systems were performed on the TACC Frontera supercomputer on nodes equipped with Intel Xeon Platinum 8280 with 56 AVX_512 logical cores per node. For the full-spike model, the 1μs simulation took ∼208,000 CPU hours; the truncated spike systems, running for 4μs, took ∼925,000 CPU hours on average.

### 7. Analysis of macromolecular spatial organization

Interactions between proteins were investigated using a computational method for analysis of relative orientations (67). In a first pass, protein representations were reduced to their center of mass positions and orientations of pre-specified internal coordinate frames. The latter were determined for the M protein’s TMD and endodomain separately in their dimeric forms, S trimer, and E pentamer by aligning the axis of symmetry with the *z* axis and aligning the vector from the center of mass of the multimer to the center of mass of the first monomer with the *x* axis. The orientation for a protein complex is then obtained by a transformation required to align the reference orientation to the actual orientation, which is stored as the translation vector and the rotation matrix. This greatly reduces the number of degrees of freedom, allowing for more efficient follow-up analyses.

In the second stage of the analysis, all pairs of protein complexes within a specific distance (10 nm) were collected, and for each pair the orientations were expressed as relative orientations by aligning one partner with the Cartesian axes, i.e., by subtracting the center of mass and multiplying by the transpose of the rotation matrix. These relative orientations were subsequently expressed as a distance and five angles, adapted from the previous work (67). For each protein, this yields an angle α, which denotes the angle of the center of mass of the partner with the XY plane, and angle β, which denotes the position of binding in the XY plane. The fifth angle captures the tilt, but is not investigated further in this work.

For all pairs of proteins within a given distance, their parameters were collected, and density plots were made to show which combinations of parameters were characteristic for each interaction. These characteristic orientations were used for assessment of the higher order structures. In addition, these orientations were combined with per-protein XY-plane particle densities to compare with the related experimental data obtained from the Cryo-EM images of SARS-CoV (8).

In addition, to eliminate the possibility that the key feature of M dimers to form the filament-like assemblies was due to the initial setup, a flat system of 41 randomly positioned M dimers was simulated for an extended period of 13μs. For this simulation, we selected the same lipid composition as model M1.

### 8. Shape analysis

To assess the shape of the SARS-CoV-2 envelope the eigenvalues of the gyration tensor as well as two shape descriptors, *S* and *Delta*, were computed following their analysis (68). Delta descriptor measures the asphericity of a particle distribution with a lower bound of 0, which indicates a perfect spherical distribution. *S* descriptor, in contrast, can have negative values indicating a point distribution corresponding to an oblate spheroid and positive values indicating a distribution corresponding to a prolate spheroid. Furthermore, to measure convergence of the simulation we estimated the auto-correlation and equilibration times as previously proposed (69, 70). In addition to this global shape analysis, a more detailed view was determined following a procedure derived from an efficient method to calculate a molecular hull and contact bodies (71), which consists of extruding a triangulated sphere to the surface of the virion, in this case marked by the M dimer TMD centers of mass. The extrusion of each vertex was determined from a kernel density estimate of TMD distances from the center in angular coordinates. This yields a smoothed triangulated surface through the membrane center, which allows visual inspection of the dynamics of the shape, and from which shape volumes and surface areas can be calculated easily.

### 9. Protein network analysis

To get further insights into the dynamic reordering of structural proteins that took place in the envelope, we conducted the connectivity analysis, constructing domain-level protein-protein interaction networks. This approach was a scaled-up version of the protein structure network (PSN) analysis, which was previously employed in the structural characterization of individual proteins (72-75). Here, we defined a network node not as an individual amino acid residue, but as an entire protein domain. In this analysis, we differentiated only between the proteins’ transmembrane domains (TMD) and endodomains (ED), which resulted in two separate physical contact networks: one within the lipid membrane and one on the inside of the virion.

We processed the reduced trajectory files (see Section 6 in Materials and Methods) that preserve the center of mass and orientation for each protein, substituting each entry with the canonical pdb model used in this study and conducting contact analysis with a cutoff distance of 0.6nm, a frequently used threshold for the coarse-grained structures (76-78). This procedure is performed separately on TMDs and EDs of the structural proteins. As a result, two sets of temporal dynamic networks for each of the truncated spike models, M1 and M3, were obtained. Each temporal dynamic network was a series of domain-domain interaction network snapshots, taken every 1ns, from 0μs to 4μs. Several key network properties were calculated for each network snapshot: average degree, number of connected components, average diameter, transitivity, and average degree connectivity for nodes with degrees 1, 2, and 3.

The degree *d*_*i*_ of a node *i* is the number of edges incident to it. The average degree in a network of *N* nodes is calculated according to the following formula: 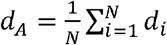. The number of connected components is the minimal number of subgraphs that have a path for any pair of vertices. The diameter of a graph *G* is the maximum length of the shortest paths for each pair of vertices (79): max_*i*_,_*j*∈*G*_ *δ*(*i, j*), where *δ*(*i, j*) is a shortest path between vertices *i* and *j*. Because our network was composed of multiple connected components, we calculated the diameter for each connected subgraph, and defined an average of the diameters: 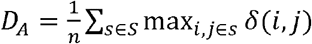, where *s* is a set of connected components and *n* is its cardinality. The transitivity (also called the clustering coefficient) is the relative number of triangles (#*triangles*) present in a given network compared to the number of all possible triangles (#*triads*): *T* = 3 (#*triangles*)/ (#*triads*) (80). Finally, the average *k*-degree connectivity is the average degree of the nearest neighbors for the nodes with the degree *k* (81). In this study, we computed the average degree connectivity for *k* =1, 2, and 3.

The number of connected components over time was fit with exponential and biexponential models:

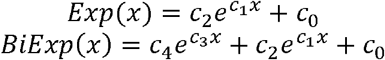

The resulting models were compared using Akaike’s information criterion (AIC) and Bayesian information criterion (BIC) (82). Results were plotted using Python 3.9 (83) and Bokeh library (84). A Sankey diagram of connected components was plotted in MATLAB (85) using visualization module of PisCES algorithm (86).

### 10. Visualization of simulation trajectories

Renderings of the virus envelope structure and MD simulation were produced using VMD (87) on TACC Frontera’s large-memory nodes with 2.1 TB NVDIMM memory. A coordinate file containing the molecular structure was used as an input for VMD. Once loaded, each coarse grain bead was represented as a sphere with a radius three times its van der Waals radius. Solvent molecules were excluded from the visualization. To capture the whole dynamic simulation process, one frame per one nanosecond was extracted from the original trajectory using *gmx trjconv* (64); the extracted XTC file containing the frames was then loaded into VMD. Lastly, a *tcl* script was used to visualize each frame, followed by rendering the frames into TGA images using tachyon (88).

Visualization of the simulation trajectory of M1 (Suppl. Movie S3) was made by rendering each frame in the 4μs trajectory of the envelope model M1 and rotating 2,000 steps around y-axis clockwise and 2,000 steps counterclockwise; each step was 0.9 degrees. Another visualization of the simulation trajectory of M1 (Suppl. Movie S4 was made with imagemagick montage (89) by showing two identical 4μs trajectories, first one revealing the outside and the second one revealing the inside of the envelope’s structural model M1. For Visualization of 1μs trajectory of model M2 (Suppl. Movie S5), we used FFmpeg to convert images into a MP4 movie with 60 fps. Rendering a movie with 4,000 frames corresponding to 4μs of simulation took ∼670 CPU-hours on TACC Frontera supercomputer.

To visualize the protein network (Suppl. Movies S7 and S8), we first convert each node’s 3D position into its 2D projection. We used the Mercator projection, which is a cylindrical projection that preserves local directions and shapes. Specifically, for each node, its 2D projection (a,b) is calculated by:

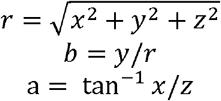

where (x,y,z) is its normalized 3D position. The orientation of each protein domain is also converted in a similar manner. Then, we plot each node as a ellipse, where the its long axis follows the orientation and an orange tip suggests the direction, and colored spike proteins as red nodes and membrane proteins as blue nodes. The edges are represented as black lines where the new edges from the previous frame are highlighted as orange. We generated 1 frame per 10 ns, which resulted in 400 frames corresponding to 4μs of simulation.

## DISCUSSION

Despite remarkable similarity of the virion structures shared between betacoronaviruses, such as SARS-CoV, MHV, and SARS-CoV-2, and the efforts to elucidate the structure of these viruses using imaging and computational methods (29, 35, 90-93), no high-resolution structure of the envelope currently exists. Our integrative approach allows combining experimental information at different resolutions into a consistent model, providing structural and functional insights beyond what can be obtained by a single experimental method. The obtained model is an important step towards our understanding of the underlying molecular architecture of the entire virus and successfully bridges the gap between molecular simulations and electron microscopy of virions, reproducing the experimentally observed density profiles of the local envelope structure. Furthermore, the developed computational protocol can be applied to study the envelopes of other coronaviruses, once the models or experimental structures and stoichiometries of the structural proteins comprising the envelope are obtained. The model will join in other efforts to structurally characterize virion particles with molecular dynamics such as influenza A, human immunodeficiency virus (HIV), and hepatitis B virus (HBV) (94-98).

The model can serve as a structural scaffold for understanding the interplay between mutual orientations of neighbor spike trimers and the role of this orientation in the viral interaction with the host receptors, as well as for studying the interactions between the proposed elongated form of M dimers and N-proteins(8) to discern the mechanistic determinants of the virion’s stability. The structures of M dimers and the complex inter-dimeric assemblies they form can provide the structural basis for understanding the molecular mechanisms behind the viral assembly. The structural knowledge of complexes formed by M proteins can also be helpful when designing new antiviral compounds targeting the interaction interface of the M protein and thus preventing formation of the envelope, an approach recently suggested for other viruses (99, 100). Targeting of the viral envelope with antiviral drugs is directly accessible within the Martini model framework (101). Finally, the structural model of the viral envelope will facilitate the development of viral-like nanoparticles for novel vaccines (102).

## Acknowledgments

We would like to thank Frank Alber, Sai Ganesan, Kuang Shen, and Celia Schiffer for the useful feedback about the manuscript. Part of computation was performed on resources from Juelich Supercomputing Centre. The authors acknowledge the Extreme Science and Engineering Discovery Environment (XSEDE) Texas Advanced Computing Center (TACC) at The University of Texas at Austin for providing HPC resources that have contributed to the research results reported within this paper. W. P. acknowledges funding from the Novo Nordisk Foundation (grant No. NNF18SA0035142) and INTERACTIONS, Marie Skłodowska-Curie grant agreement No 847523.

## Author contributions

All authors contributed extensively to the work presented in this paper. DK jointly conceived the study with SJM. WP, FG, ON, SL, TAW, and DK developed the methodology. WP, FG, ON, SL, and TAW performed the experiments and collected data. WP, FG, ON, SL, TAW, SJM, and DK analyzed data. WP, FG, ON, SL, TAW, and DK generated visualizations for different parts of the project. ON, SL, WP, TAW, and DK wrote the original draft, and all authors contributed to the review and editing of the manuscript. SJM and DK supervised and managed the project.

