## Supplementary Information for "Molecular architecture and dynamics of SARS-CoV-2 envelope by integrative modeling"

**Lipid fingerprint analysis**

Membrane proteins are known to collect certain lipids in their immediate lipid shell. Analyzing this lipid environment preference is referred to as lipid fingerprinting [1]. Here, we have conducted the lipid fingerprinting analysis of the envelope proteins under consideration of clustering of M dimers in a model of the lipid membrane that mimics composition of the endoplasmic reticulum (ER). Based on the results, we then slightly adjusted the lipid composition of the viral envelope following the reasoning that the high concentration of M dimers likely leads to the enrichment of lipids preferentially found in the M dimer lipid shell.

The molecular dynamics simulation of M dimers in the flat membranes also showed their tendency to cluster (Supp Fig. S11), thus the lipid fingerprint was extracted from simulations with four M dimers after they formed clusters in the modelled membrane. For E and S proteins, one corresponding homooligomer was added to the four M dimers setup. In addition, a simulation with a single M dimer was run to see if the results were different. The second factor that we investigated was the influence of cardiolipin on the lipid fingerprint. For the ER composition that contained cardiolipin, the charge state of cardiolipin was not specified. The charge state can either be neutral, singly, or doubly charged. Here, we used the doubly charged version, to see if there was a strong effect on the lipid fingerprint. In addition, we ran a set of simulations without cardiolipin. For all systems, the lipid fingerprint was analyzed per a single oligomer, but averaged over the oligomer copies in the cluster, accounting for the clustering properties of the M dimer.

In the depletion enrichment analysis for the M dimer system, the overall trend showed slight depletion of cholesterol and POPE independent of a monomeric or oligomeric state of M protein, or whether it was in a cluster (Supp. Table S1). POPC was neither enriched nor depleted. POPS, (a charged lipid) was slightly enriched independent of the oligomeric state or clustering. Interestingly POPI, also a charged lipid, was enriched in the simulations without cardiolipin, but depleted in the simulations with cardiolipin, where cardiolipin was strongly enriched. A possible reason could be that the doubly charged cardiolipin had a stronger affinity for the M dimer and displaced POPI in those simulations. This sequestering of negatively charged lipids was consistent with recent results of all-atom simulations of isolated M-dimers [2]. Our findings were further confirmed by computing the relative lipid densities around the proteins (Supp. Fig. S7). Specifically, the enrichment of POPS and Cardiolipin can be clearly observed, while one could also see the absence of POPI from the regions where cardiolipin bound. The lipid densities appeared to be clearly asymmetric which reflected the asymmetry of the M-dimer itself and which was also consistent with the recent all-atom simulations [2].

The simulation of the system that included one S trimer followed the same general trend as in the case of M dimer (Supp. Fig. S7), with slight deviations in depletion and enrichment values. Interestingly, POPS and POPC were more enriched at the expense of cardiolipin and POPI. Furthermore, the system that included E pentamer exhibited similar behavior as well, however, in this case POPI was enriched even in the simulation with cardiolipin, probably at the expense of POPC that was slightly depleted (Supp. Fig. S7).

Finally we note that in spite of clear trends, we did not see strong affinity of any of the proteins for any lipid. Thus, we decided to only slightly modify the final lipid composition of the envelope based on the depletion enrichment values for the four M-dimer copies. In addition, considering the uncertainty around the charged state of cardiolipin, we decided to increase the percentage of cardiolipin, but not as strongly as suggested by the depletion enrichment analysis. This led to the final lipid composition (Table 1).

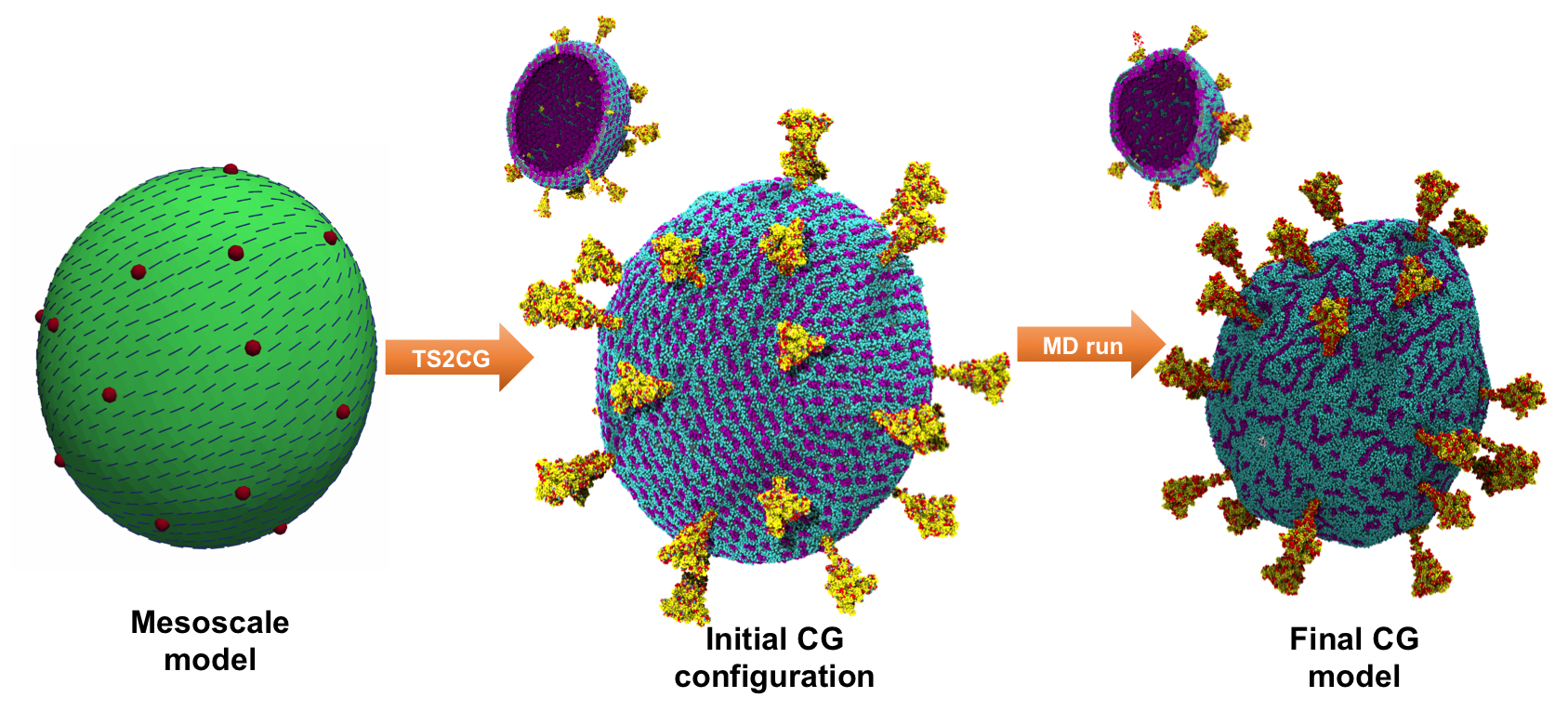

Figure S1:

Key molecular representation stages during the integrative modeling of the whole virus.

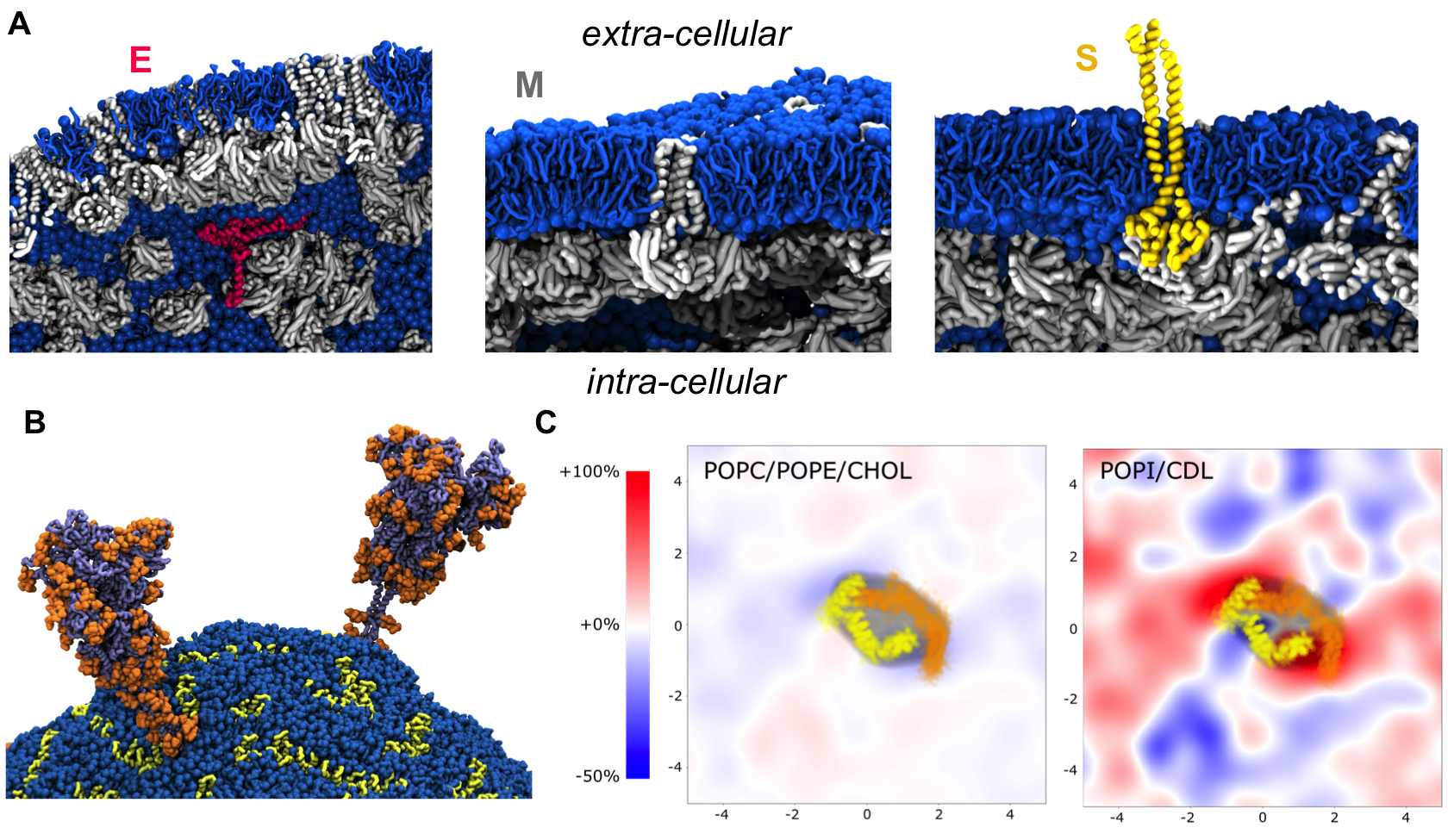

**Figure S2. Protein-lipid interactions in the envelop model:** **A.** Local view of the molecular interactions between lipids (sapphire blue) and structural proteins: E pentamer (ruby red), M dimer (silver), and truncated S trimer (gold). Unlike the E pentamer that remained fully surrounded by the lipid molecules, every S trimer became supported through molecular interaction at least by one M dimer at the end of the simulation. Similarly, most M dimers were found interacting with other M dimers. **B**. Two glycosylated S trimers (dark grey) shown together with their glycans (orange) after the 1µs simulation run. **C**. Lipid fingerprinting analysis showed a slight enrichment of the anionic lipids POPI and CDL around M dimers.

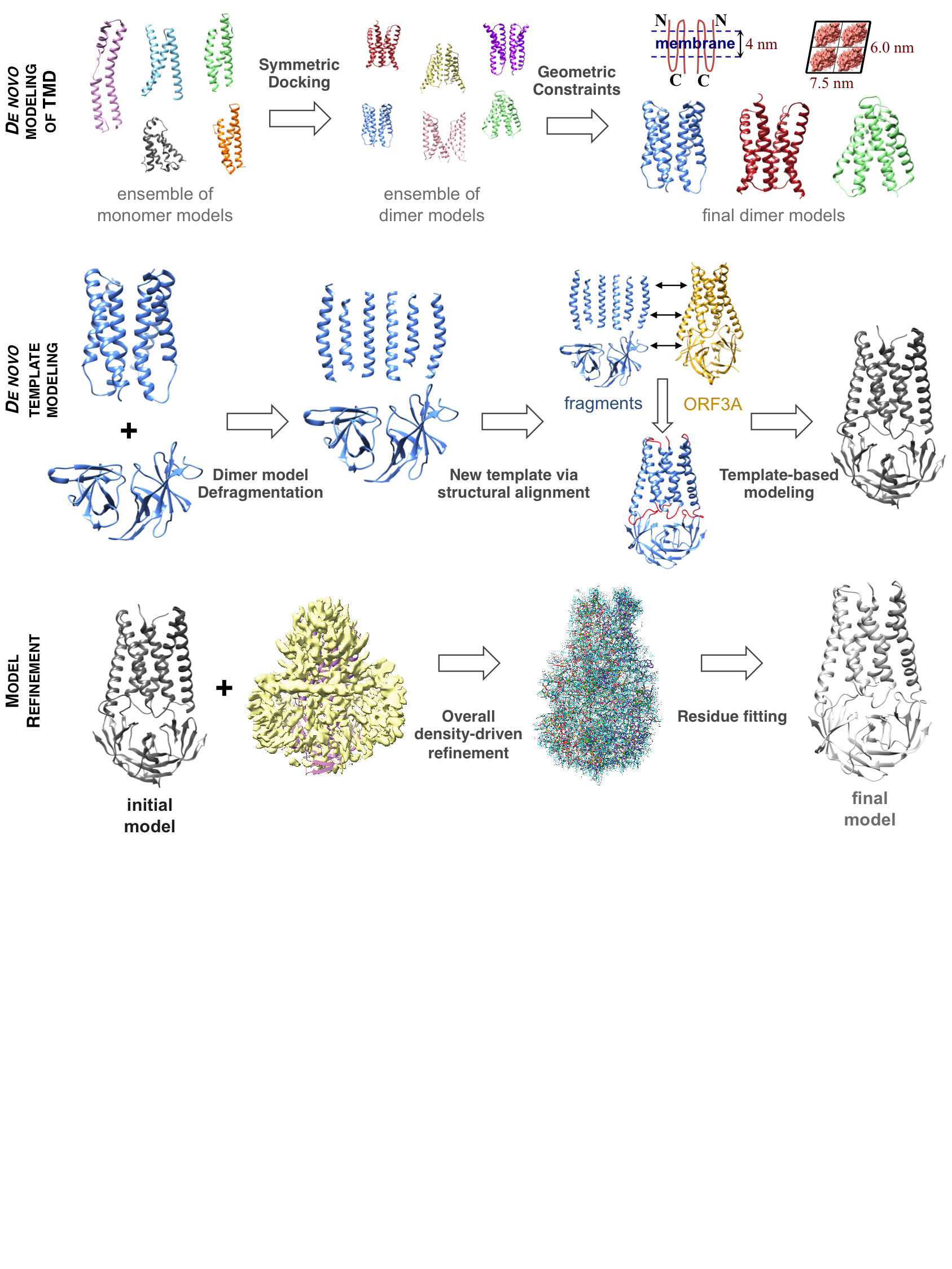

Figure S3:

Basic stages of structural characterization of M protein’s dimeric complex using integrative modeling.

**A**

```
ORF3A        1 MDLFMRIFTIGTVTLKQGEIKDATPSDFVRATATIPIQASLPFGWLIVGVALLAVFQSAS
M_Protein    1 MAD-----SNGTITV--------------EELKKLLEQWNLVIGFLFLTWICLLQFAYAN
consensus    1 *  ...... **.*...............     .  *  *  *.* .    *  *  *

ORF3A       61 --KIITLKKRWQLALSKGVHFVCNLLLLFVTVYSHLLLVAAGLEAPFLYLYALVYFLQSI
M_Protein   42 RNRFLYIIKLIFLWLLWPVTLACFVL---AAVY-RINWITGGIAIAMACLVGLMWLSYFI
consensus   61 ... . . *   * *   *   * .*...  **...  . .*.      * .*..    *

ORF3A      119 NFVRIIMRLWLCWKCRSKNPLLYDANYFLCWHTNCYDYCIPYNSVTSSIVITSGDGTTSP
M_Protein   98 ASFRLFARTRSMWSFNPETNILLN-------------------------VPLHGTILTRP
consensus  121    *.  *    *       .*  .........................*   *   * *

ORF3A      179 ISEHDYQIGGYTEKWESGV----------KDC---VVLHSYFTSDYYQLYSTQ-LSTDTG
M_Protein  133 LLESELVIGAVILRGHLRIAGHHLGRCDIKDLPKEITVATSRTLSYYKLGASQRVAGDSG
consensus  181 . * .  **.   .    ...........** .... . .  *  ** *  .*..  *.*

ORF3A      225 VEHVTFFIYNKIVDEPEEHVQIHTIDGSSGVVNPVMEPIYDEPTTTTSVPLSNSLEVLFQ
M_Protein  193 ------------------------------FAAYSRYRIGNYKLNTDHSSSSDNIALLVQ
consensus  241 ..............................        *      *     *  . .* *
```

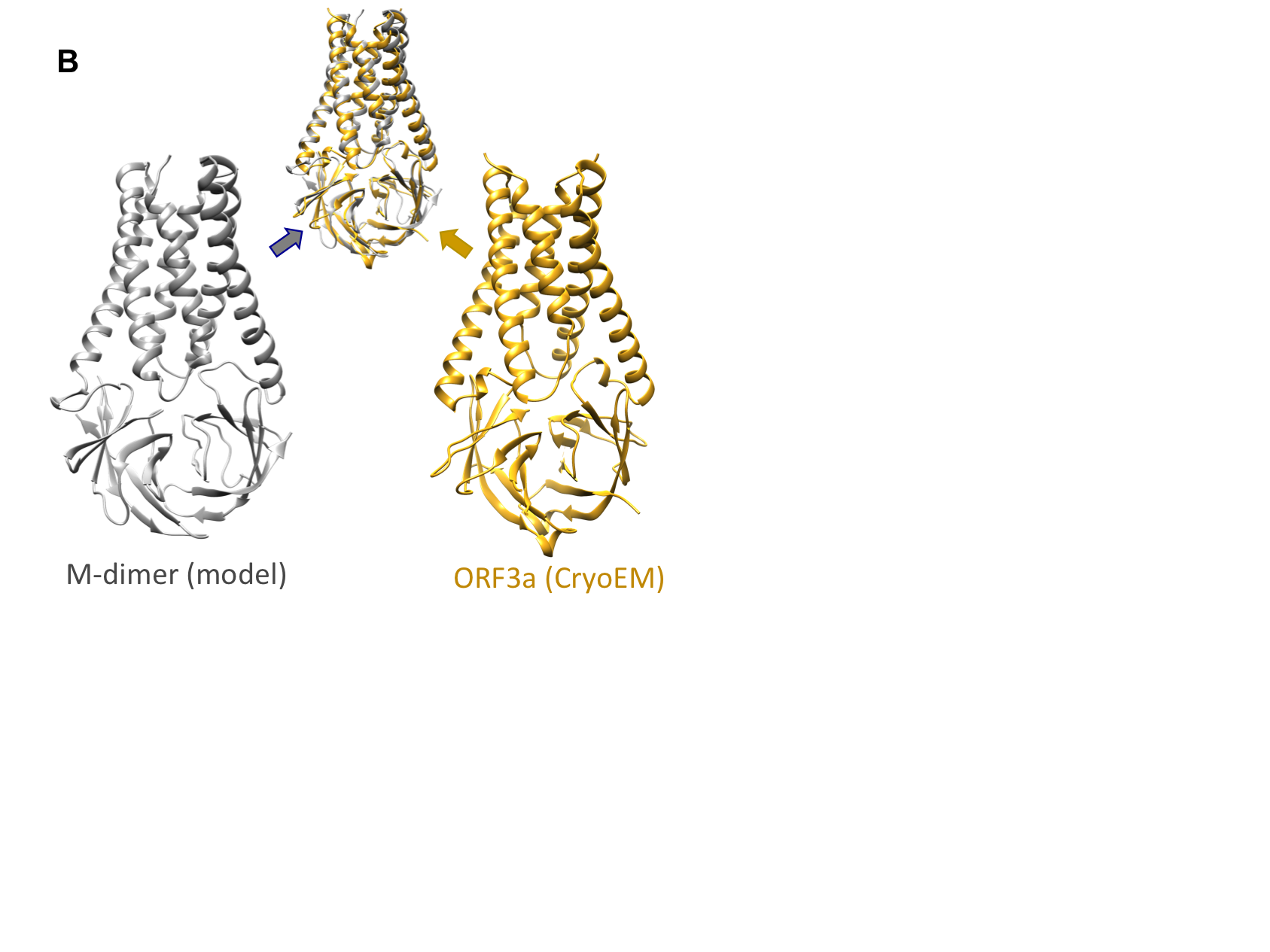

Figure S4:

In spite of substantial difference in protein sequences M model resembles closely a structurally resolved ORF3a dimer: **A.** Protein sequence alignment between M and ORF3A proteins of SARS-CoV-2 using TCoffee. Sequence identity is 16%. **B:** Structural alignment of M dimer model and ORF3A structure (PDB ID: 6XDC) using TM-align. All-residue RMSD is 1.64A.

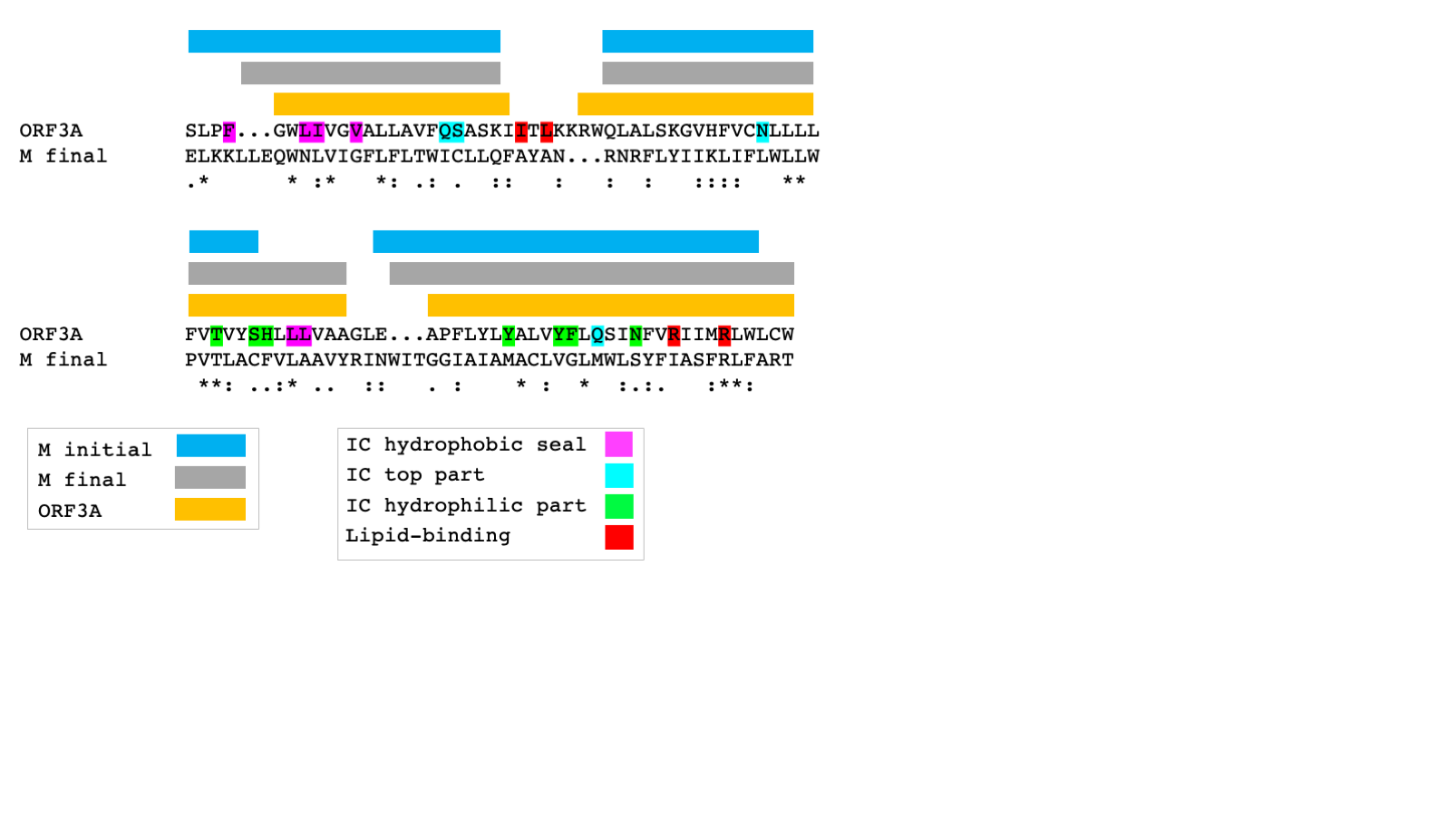

Figure S5:

Three alpha helices assigned for initial and final models of M as well as for the structures of ORF3A. The secondary structure of the final model is better aligned with the secondary structure of ORF3A. Shown is a sequence alignment of the final model of M and ORF3A derived based on their structural superposition without any sequence-based alignment optimization. The functional annotation of ORF3A show that many positions that either correspond to the functional residues or are in their immediate structural vicinity remain conserved.

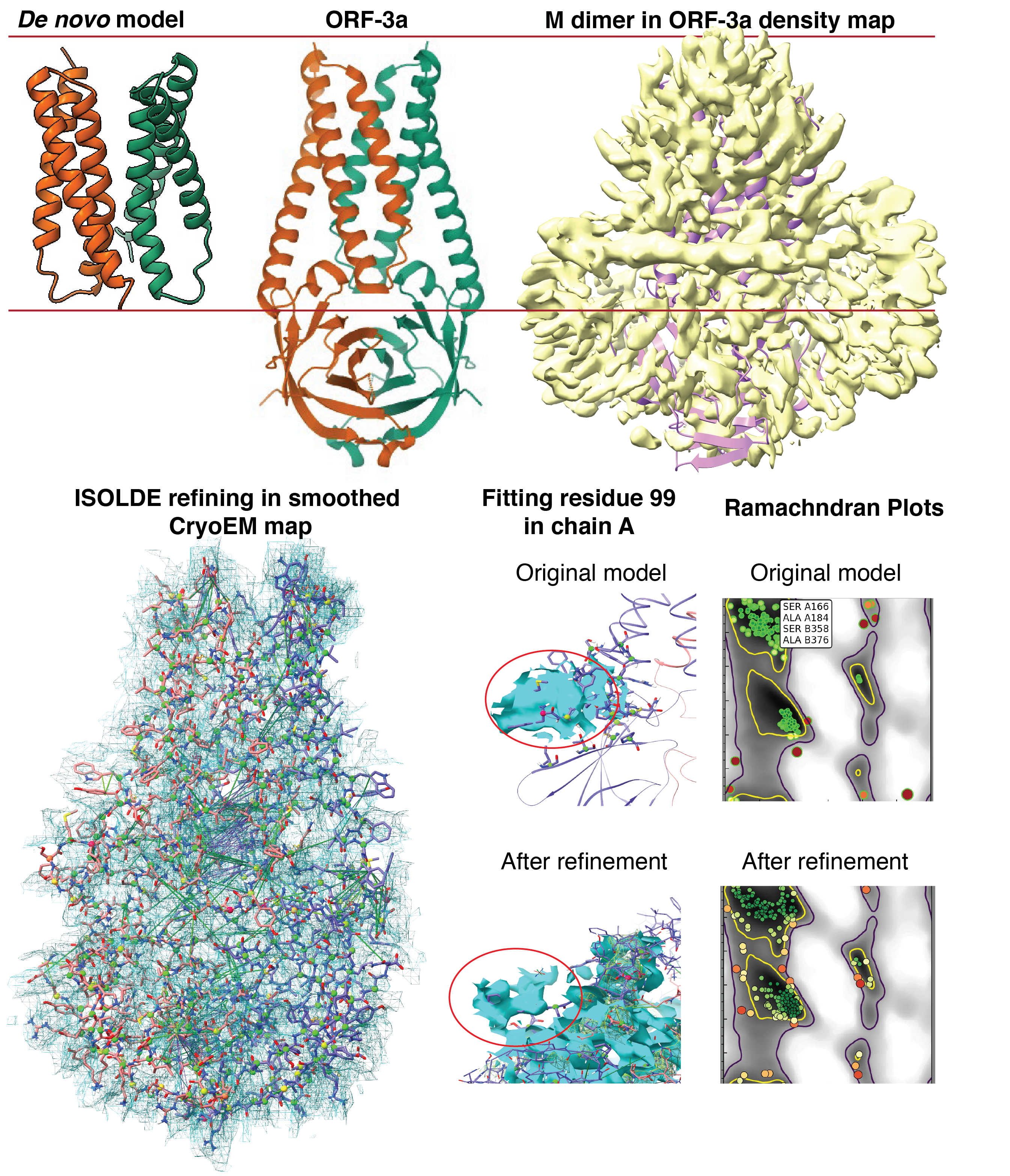

Figure S6:

Structural refinement of M dimer. Top left: *de novo* model of the M dimer without endodomain exhibit similarity to the ORF-3a protein from SARS-CoV-2. Top right: M dimer model with endodomain in Cryo-EM density map of ORF-3a in lipid nanodisc (EMD-22139). Bottom left: M dimer model during ISOLDE simulation run in ChimeraX molecular visualization suite; blue mesh corresponds to the CryoEM map with Gaussian smoothing with B-factor equal to 100, green lines correspond to the local restraints at the distances of 1.5Å or more, and the colors of residue’s atoms correspond to atoms’ goodness of fit. Bottom middle: an example of fitting a specific residue 99 in chain A; in the original model this residue sticks out of the density map (blue surface), and during the process of refinement it acquires a better fit. Bottom right: Ramachandran plots of the original and refined models; one can see that the original model contains multiple outliers (dark red), compared to the model after refinement, which could pose severe problems during molecular dynamics simulation run.

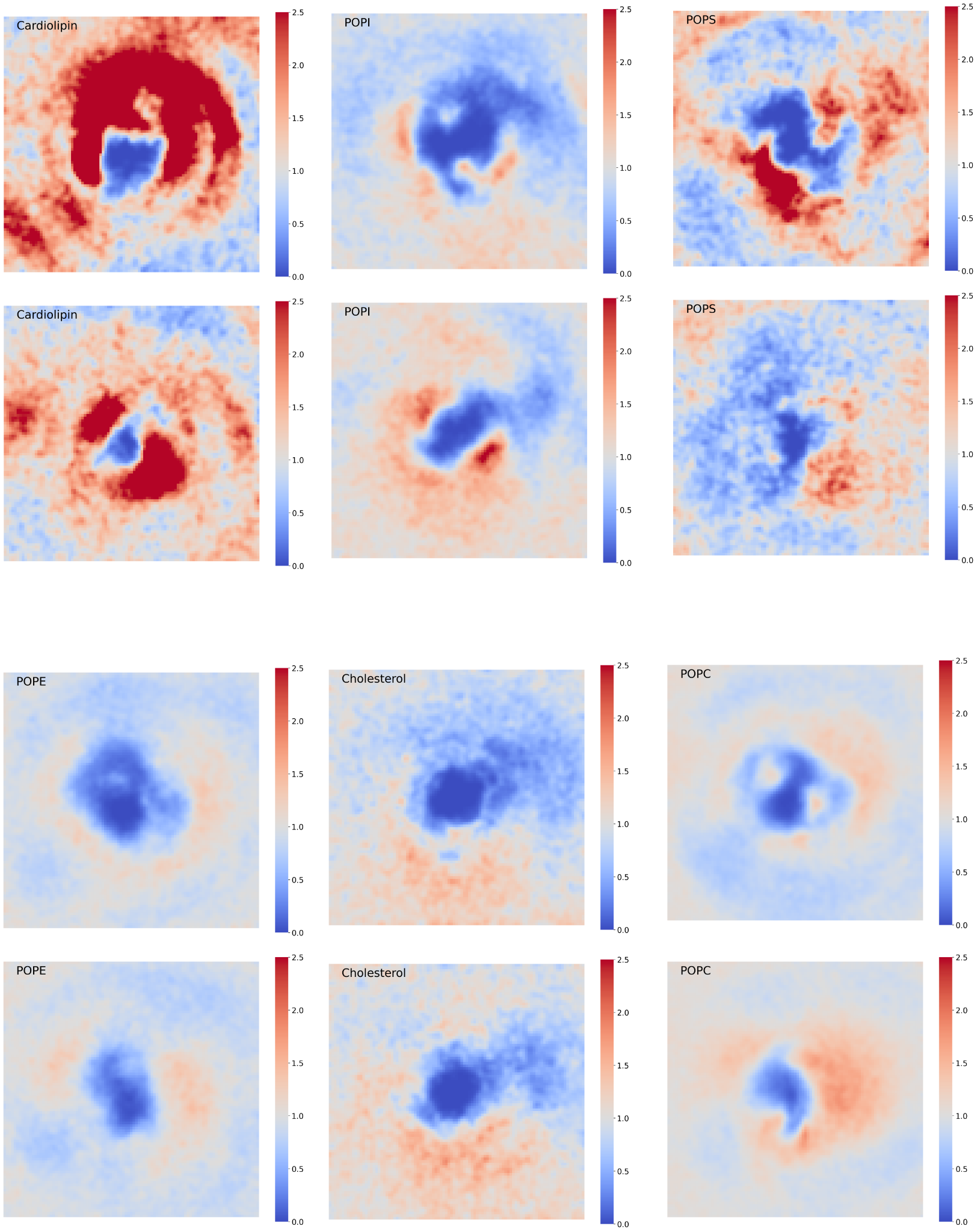
**Figure S7:**

Relative lipid densities around clusters of M-dimers. The scale goes from 0.0 for completely absent to 2.5, which corresponds to 2.5 fold enrichment over the bulk lipid density in that bin. Upper panels correspond to the outer leaflet and lower panels to the inner leaflet. Proteins are located in the center of the plot and characterized by the absence of lipids.

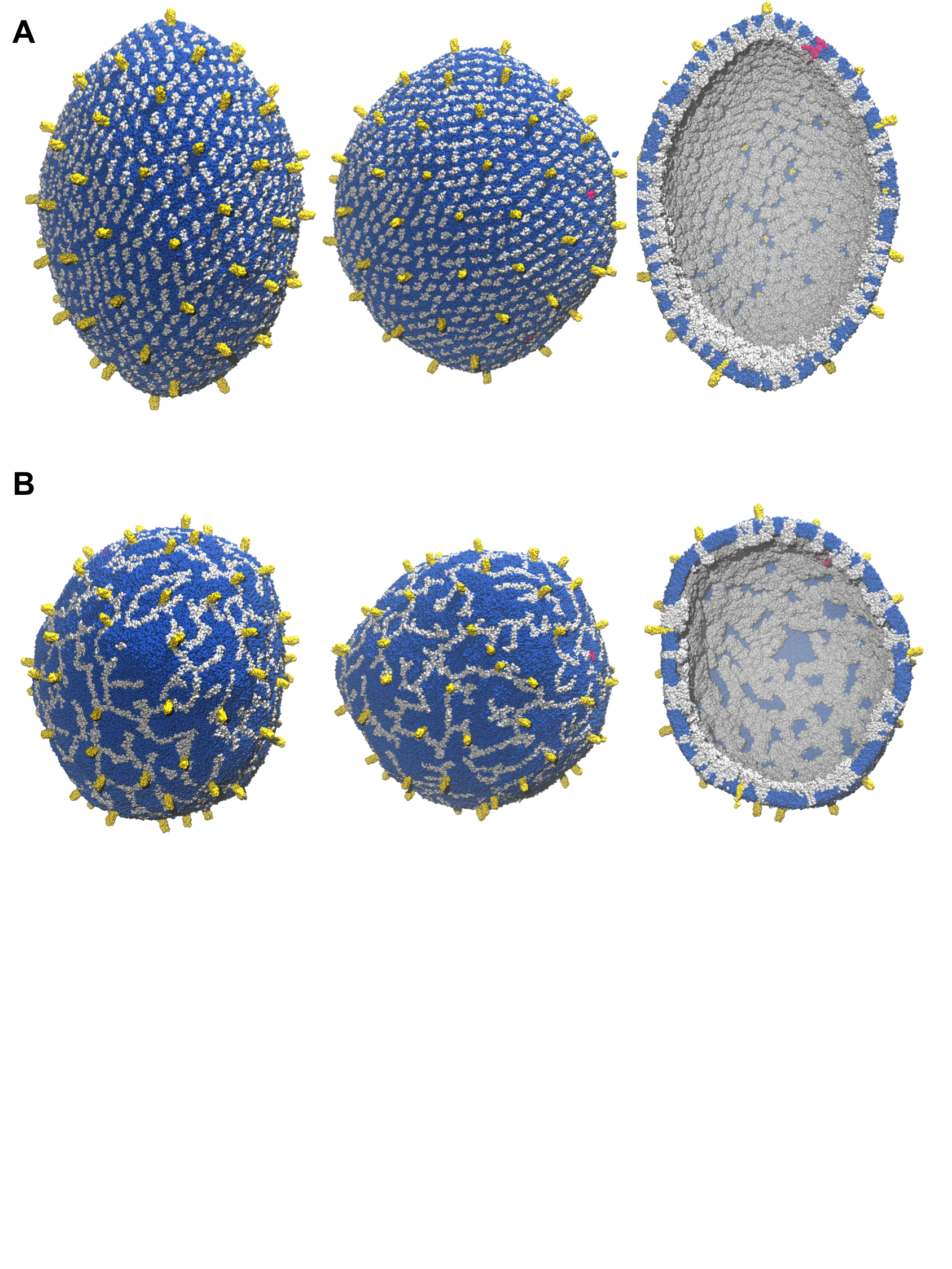

Figure S8:

Structural characterization of model M3 generated from molecular composition C2 (3 E pentamers, 71 S trimers, and 1080 M dimers) during 4 μs simulation. Lipid molecules are depicted in sapphire blue, E pentamers in ruby red, M dimers in silver, and S trimers in gold. **А:** Representation of the model at T = 0 μs. **B:** Representation of the model at T = 4 μs. Shown for each model are side view (first column), top view (second column), and inside of the envelope (third column).

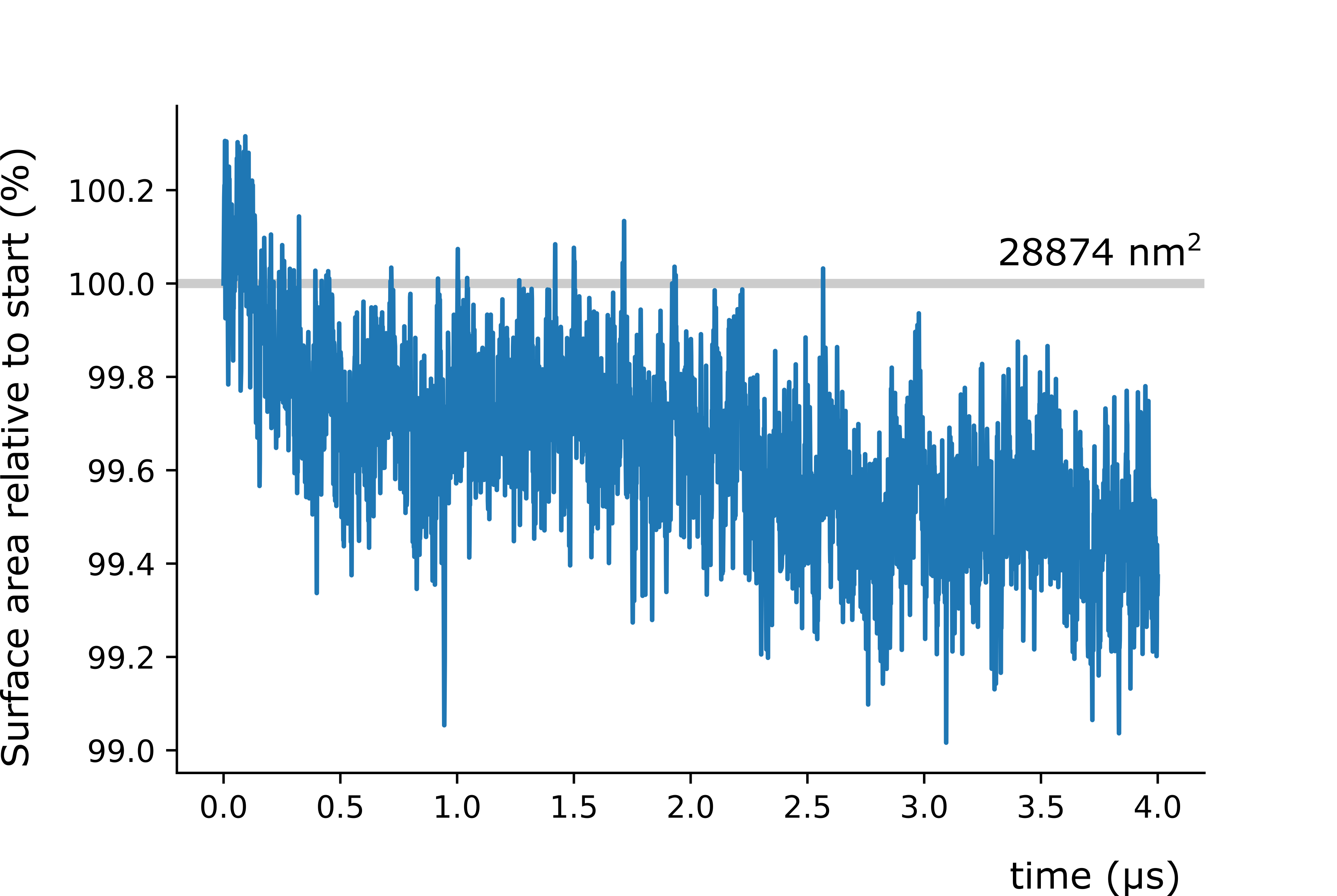

Figure S9:

Change of the envelope’s surface area throughout simulation for model M1 obtained from molecular composition C1 (2 E pentamers, 25 S trimers, and 1,003 M dimers).

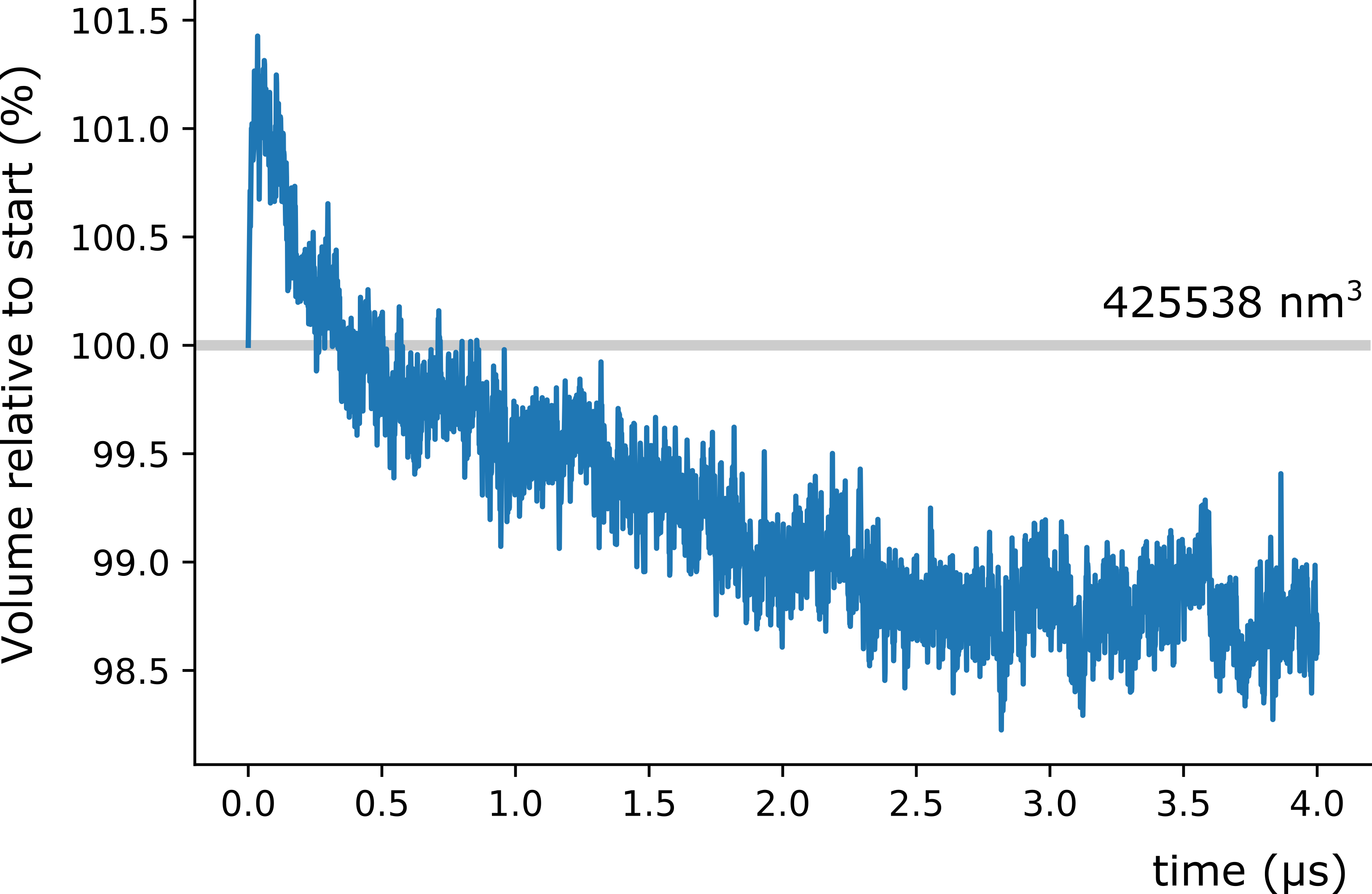

Figure S10:

Change of the envelope’s volume for molecular composition C1 (model M1) throughout simulation.

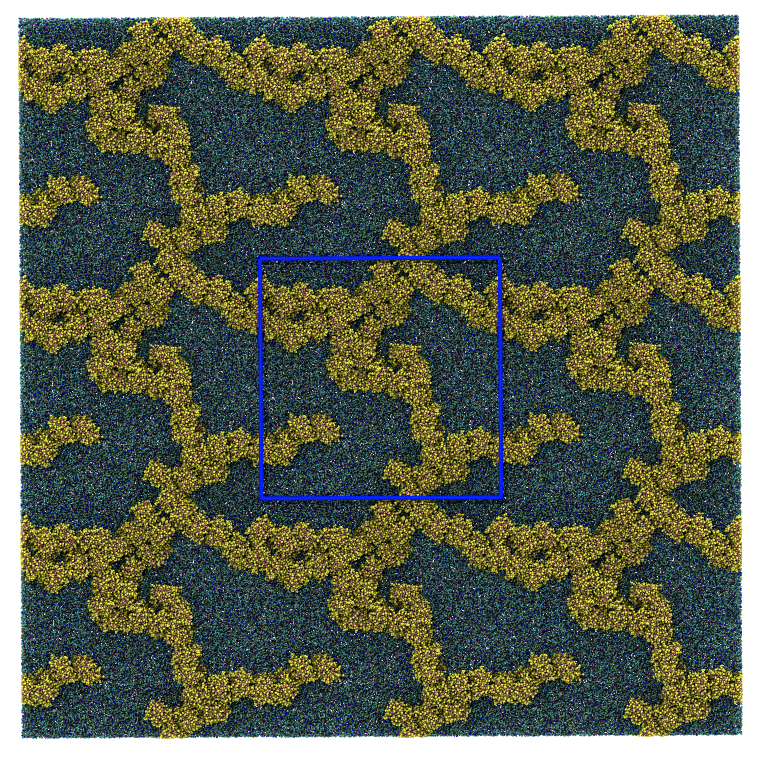

Figure S11:

13 microsecond simulations of 41 M dimers in a flat bilayer containing POPC 948, POPE 210, POPI 484, CHOL 252, POPS 196 in the upper monolayer and POPC 831, POPE 184, POPI 425, CHOL 221, POPS 167 in the inner monolayer. The pattern formed by M dimers resembles the patterns as observed in the full envelope simulations. The dynamic evolution of this system is shown in Movie S6.

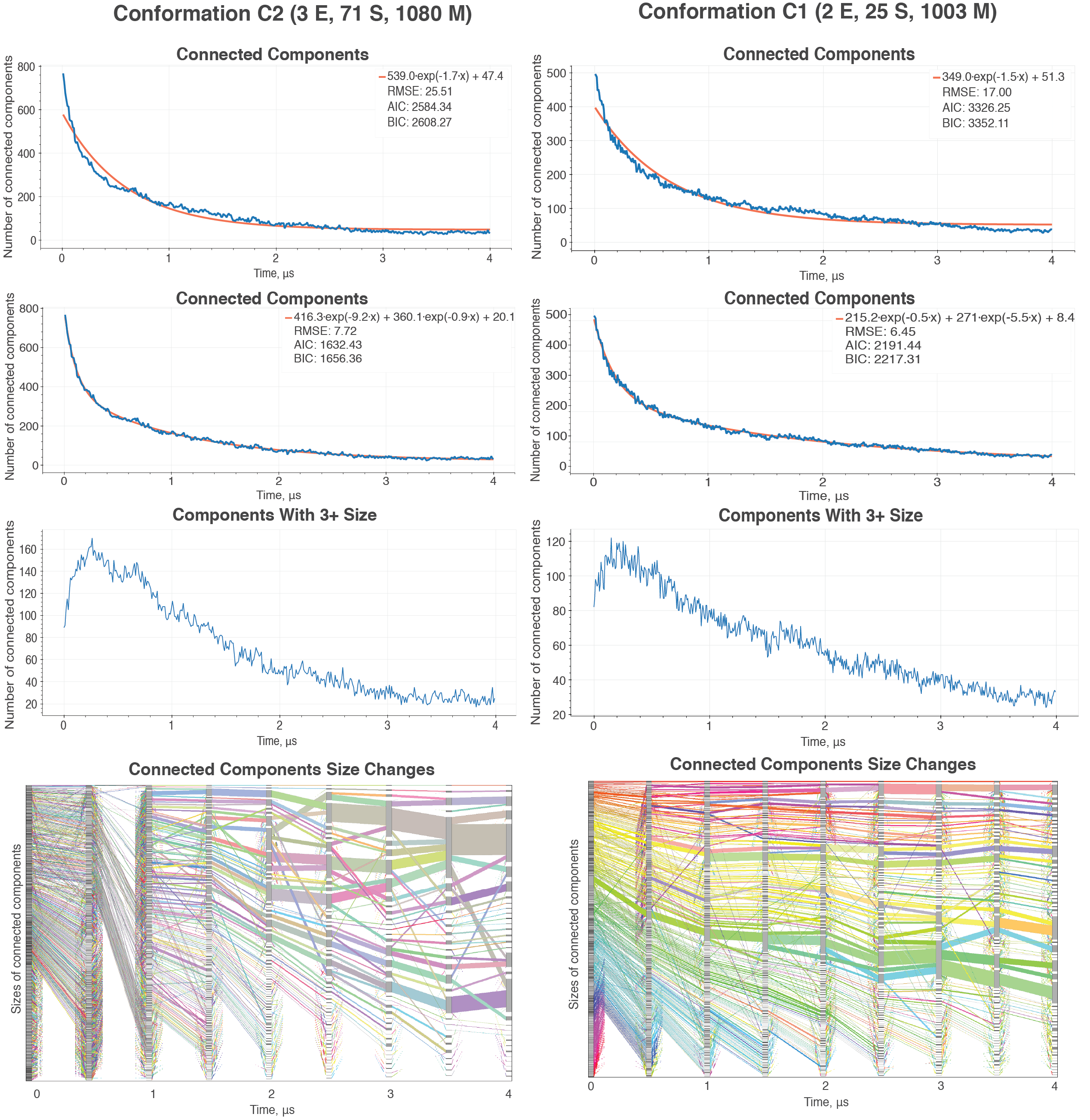

Figure S12.

Change in the number of connected components in the TM domain-domain interaction networks throughout the simulation for models M3 (conformation C2) and M1 (conformation C1). Between two possible fitted lines, exponential model (first row) and biexponential model (second row), one can see that the biexponential model provides better RMSE score along with smaller values for Akaike’s Information Criterion (AIK) and Bayesian Information Criterion (BIC), suggesting that the biexponential model better represents the underlying process and supporting our hypothesis about two separate processes happening during the simulation. Specifically, for both M1 and M3 models, there are two distinct exponential processes, fast and slow, differing in speed ~10x times. Shown in the third row is the number of components with at least 3 proteins. Again, there is an increase in the number of components at the beginning of simulation, but later the components tend to merge, bringing a total number of components under 40 for both M1 and M3. The last row represents Sankey diagram of connected components, demonstrating the tendency of the envelope structure to form larger clusters of TMDs for M dimers. Each flow depicts rearrangement of 25 or more proteins.

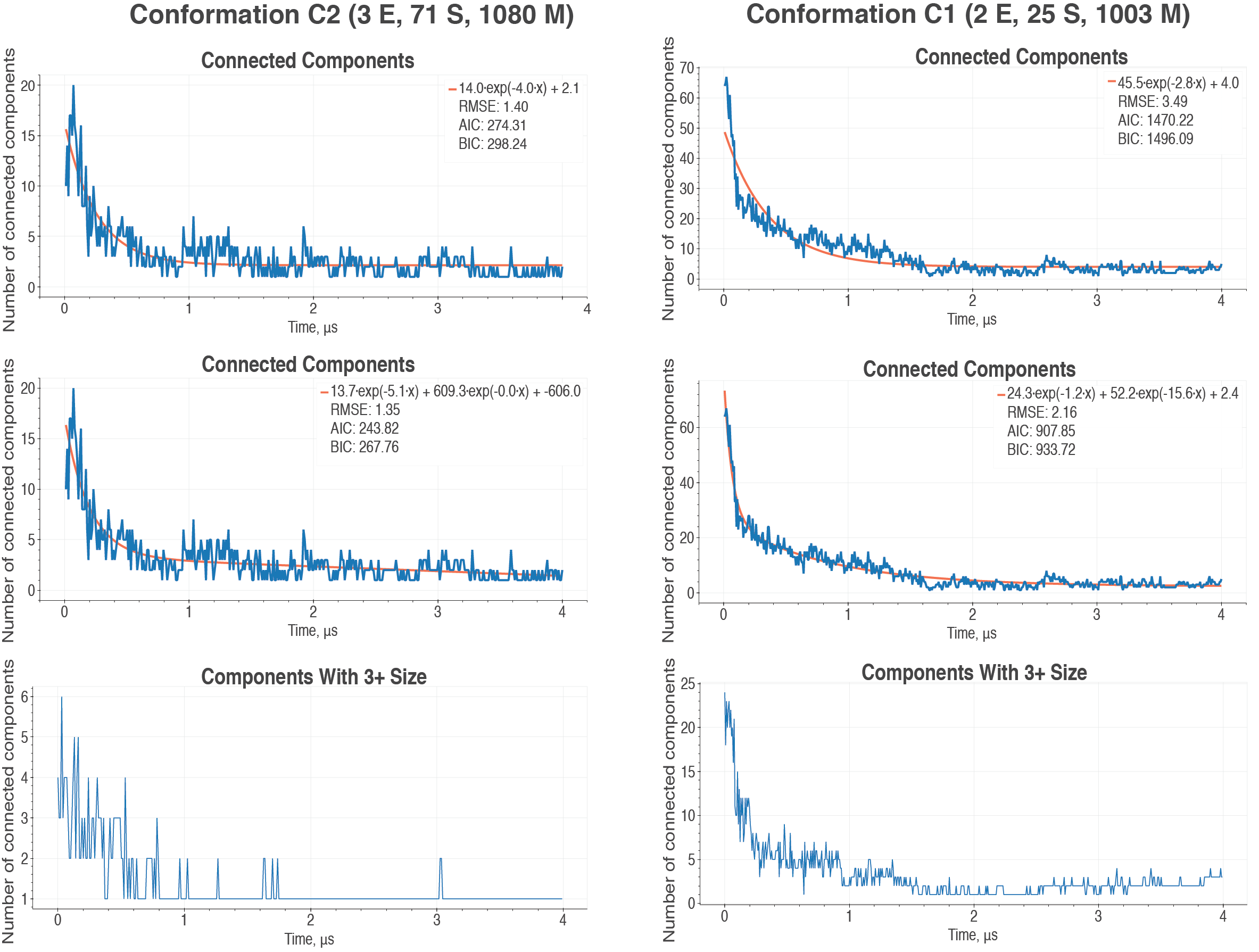

Figure S13:

Change in the number of connected components in the ED domain-domain interaction networks throughout the simulation for models M3 (conformation C2) and M1 (conformation C1). Similar to Suppl. Fig. S9, one can see that ED rearrangement can be described as a biexponential process, although for M1 the difference between biexponential and exponential models is minimal because of extremely tight packing. Also, for M3 its EDs tend to conglomerate to such an extent that they have only one, giant, connected component of the size of three or more; for M1 this number varies over time but stays under five connected components.

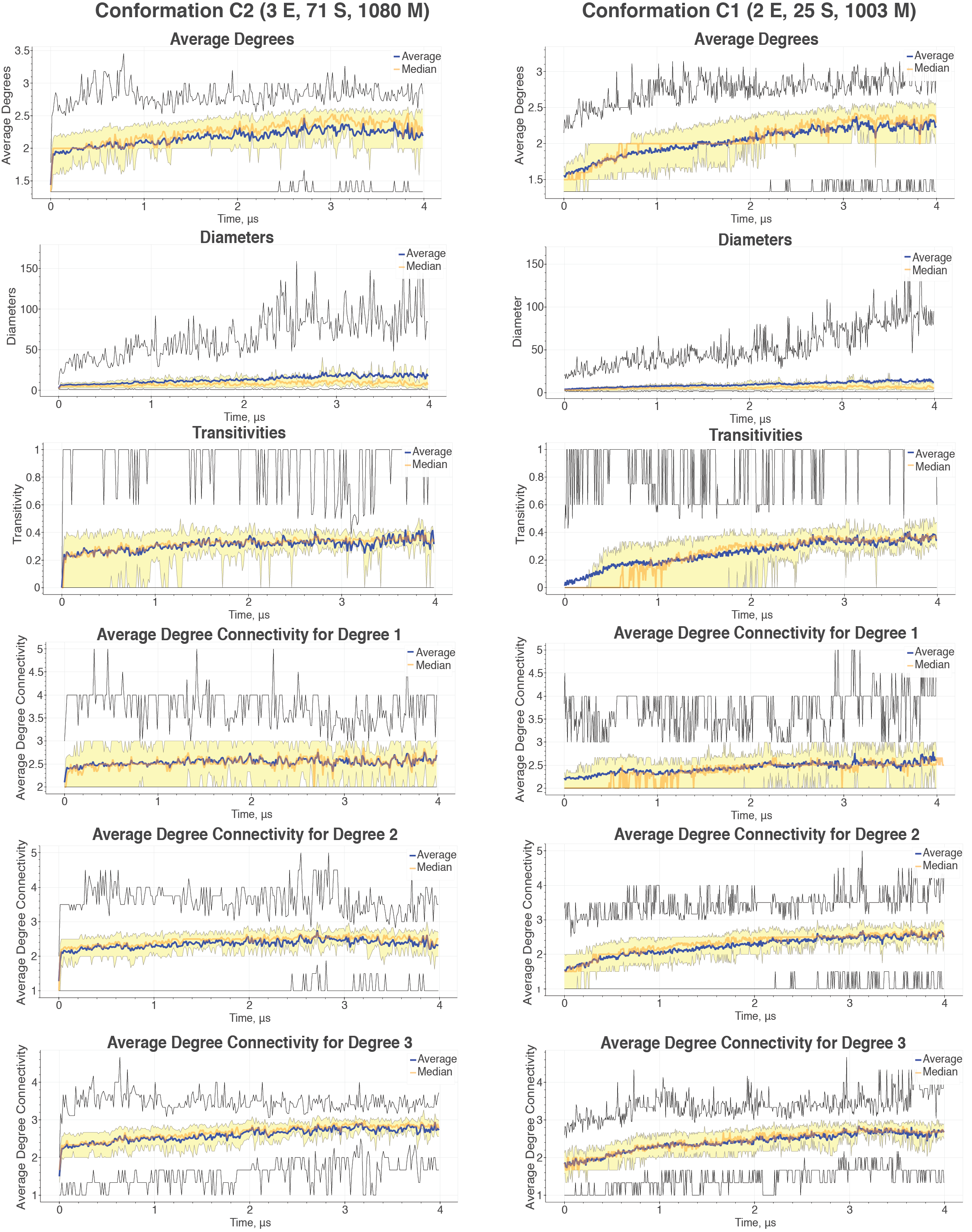

Figure S14:

Dynamics of the basic network parameters for TMD domain-domain interaction networks during the simulation of models M3 (conformation C2) and M1 (conformation C1). Yellow bands indicate values for the 2^nd^ and 3^rd^ quantiles, black lines denote the minimum and maximum values. There is a clear trend for the increase in node degrees for TMD components, plateauing during 3-4 µs around the value of 2.3. The diameter sizes tend to grow over time. The vast difference between the maximum and average or median values suggests that small number of connected components cover most of the network. The transitivity measure depicts a ratio of fully connected components in the network with respect to the maximum number of possible components for the given network. The measure stabilizes over time around the value 0.35 with very limited spread for the 2^nd^ and 3^rd^ quantiles, suggesting that this characteristic is similar for the most connected components. Average degree connectivity characterizes the nearest neighbors of the nodes of degree $k$. The values for the nodes of degrees 1, 2, and 3 all converge to comparable average degree connectivities (around 2.5), suggesting that those nodes are evenly distributed in the protein connectivity network.

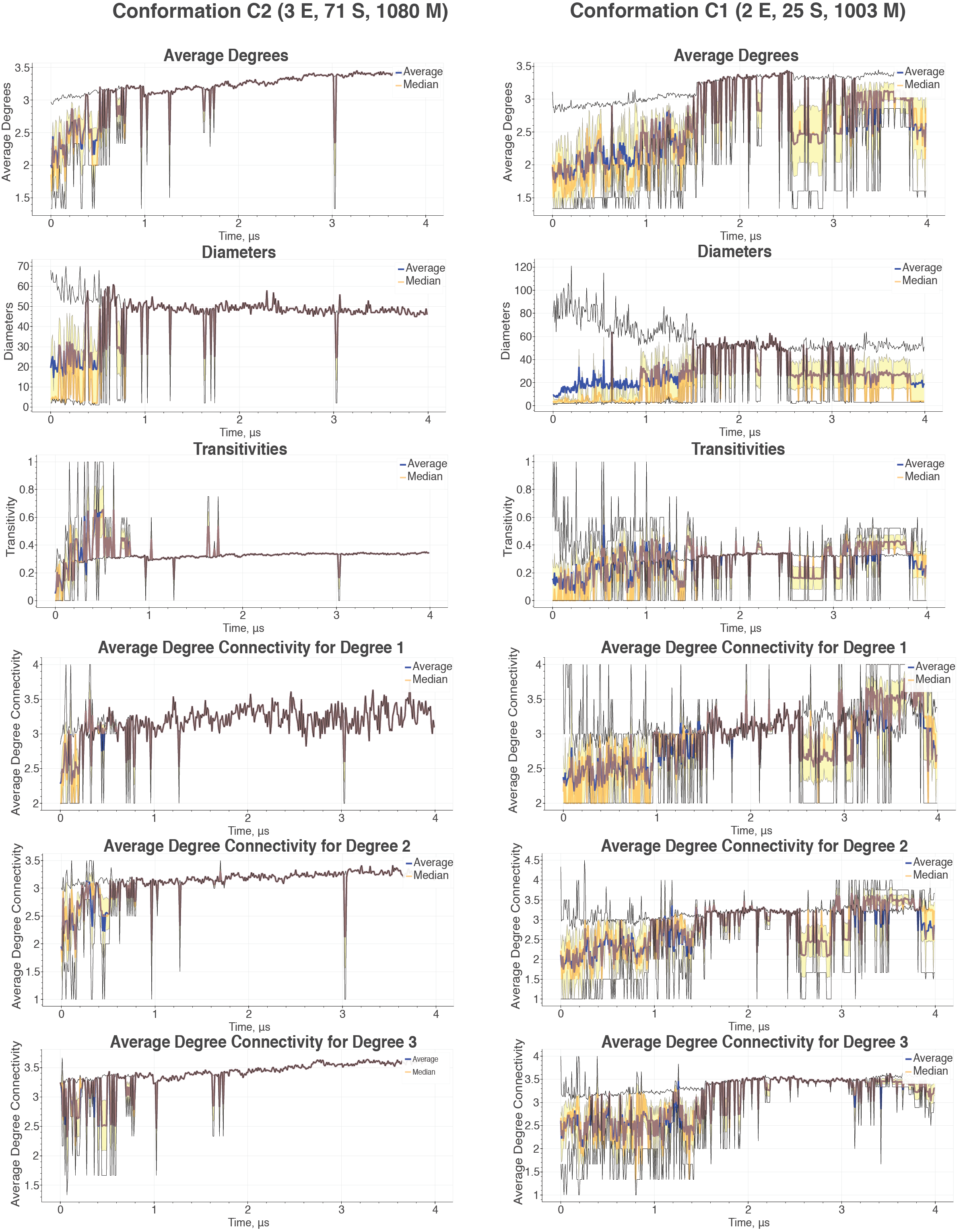

Figure S15:

Dynamics of the basic network parameters for ED domain-domain interaction networks during the simulation of models M3 (conformation C2) and M1 (conformation C1). Yellow bands indicate values for the 2^nd^ and 3^rd^ quantiles, black lines denote the minimum and maximum values. ED domain-domain network is almost fully connected, and therefore has a much smaller number of connected components. The node degrees converge to the values of 2.5 for M3 and slightly above 3 for M1. Diameters for the connected components of M3 stays around 50, while for M3 it fluctuates around the value of 30. Transitivity value approaches 0.4, which seems to be a limit for the triangulated mesh. Average degree connectivity suggests a much tighter coupling between ED domain-domain network compared with that one of TMD network.

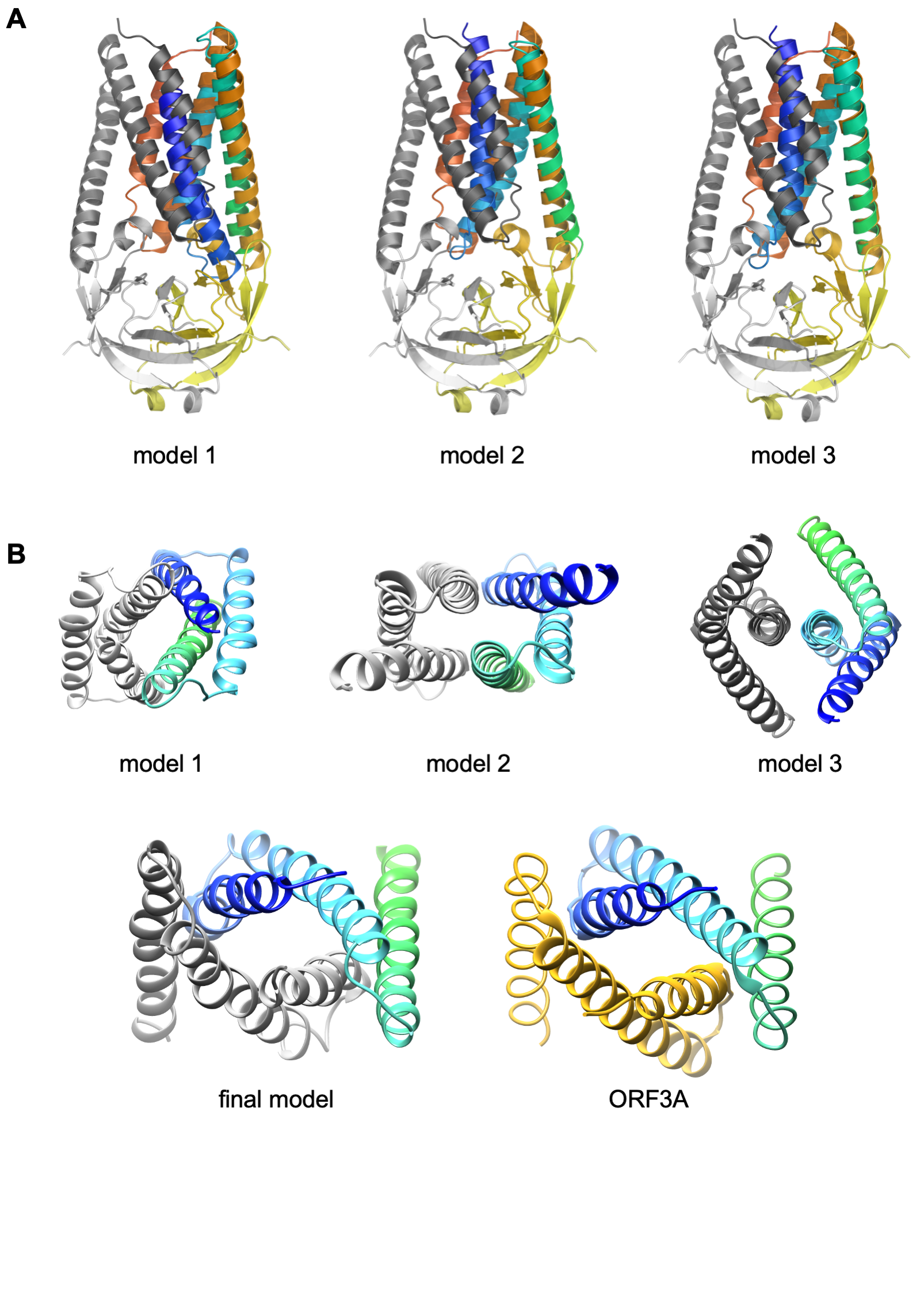

Figure S16:

Difference in the arrangement of the secondary structure elements in the transmembrane (TM) domain of M *de* novo models and the model obtained using our integrative modeling pipeline. **A:** Three initial top-scoring models of M monomer (shown only its TM domain) structurally aligned against ORF3a dimer. ORF3a has one chain shown in red-to-yellow colors, changing from N- to C-termini, and the other chain colored in grey. The M monomer is shown in blue-to-green colors, changing from N- to C-termini. While the second and third helices of M monomer (light blue-to-green) are superposed with the corresponding helices of ORF3a chain light red-to orange, the first helix of TM of M (dark blue) is shown to align with the first helix of TM of ORF3A located in the other chain (grey). **B**: Top view of three initial M dimer models (top) as compared against the integrative model of M dimer (bottom left) and ORF3A (bottom right). For all five structures one chain is colored grey (M) or golden (ORF3A), while another chain is colored dark blue-to green.

Table S1.

Depletion-enrichment index per lipid type for the three membrane proteins (M, S, E). Values are reported per multimer and averaged over copies present in the simulation. The last two rows correspond to a system without cardiolipin. Red numbers indicate enrichment while blue ones indicate depletion.

| Copies | Target | POPC (netral) | POPE  (neutral) | POPI  (-1) | POPS  (-1) | CHOL  (neutral) | CLD2  (-2) |
| --- | --- | --- | --- | --- | --- | --- | --- |
| 4M | M | 1.0063 | 0.7301 | 0.9007 | 1.0921 | 0.6995 | 3.754 |
| 4M, 1S | S | 1.1357 | 0.6643 | 0.6954 | 1.4457 | 0.6409 | 1.98625 |
| 4M, 1E | E | 0.9582 | 0.8326 | 1.1666 | 1.3291 | 0.7266 | 1.9797 |
| 1M | M | 1.0557 | 0.5283 | 1.1876 | 1.11 | 0.7601 | n/a |
| 4M | M | 0.9742 | 0.7317 | 1.1023 | 1.5782 | 0.7054 | n/a |

**Table S2.**

Number of Martini3 particles per element. Lipid molecules include 1-palmitoyl-2-oleoylphosphatidylcholine (POPC), 1-palmitoyl-2-oleoyl-sn-glycero-3-phospho-ethanolamine (POPE), 1-palmitoyl-2-oleoyl-sn-glycero-3-phosphoethanolamine (POPI), 1-palmitoyl-2-oleoyl-sn-glycero-3-phospho-L-serine (POPS), cholesterol (CHOL), and cardiolipin (CDL2). S-TM corresponds to the transmembrane domains of S trimer, S-truncated corresponds to the truncated model of S timer without its endodomains, S-full corresponds to a model of entire S trimer.

| **Element** | **N of Martini3 particles** |
| --- | --- |
| ***Protein homo-oligomers*** | |
| S-TM trimer | 459 |
| S-truncated trimer | 711 |
| S-full trimer | 3,822 |
| E pentamer | 880 |
| M dimer | 946 |
| ***Lipids*** | |
| POPC | 12 |
| POPE | 12 |
| POPI | 14 |
| POPS | 12 |
| CHOL | 8 |
| CDL2 | 27 |

**Table S3.**

Overview of an alternative system composition. Shown are the compositional details of a stable envelope model for molecular composition C2 (3 E pentamers, 71 S trimers, and 1080 M dimers) with truncated (short) S trimers. A CG water particle corresponds to 4 real water molecules.

| **Composition** |  | C2, 4μs |
| --- | --- | --- |
| **Proteins** | S | 71 |
|  | E | 3 |
|  | M | 1,080 |
|  | **Total particles** | **1,056,909** |
| **Lipids** | POPC | 39,755 |
|  | POPE | 13,191 |
|  | POPI | 7,195 |
|  | POPS | 0 |
|  | CHOL | 0 |
|  | CDL2 | 0 |
|  | **Total particles** | **736,082** |
| **Solvent** | Na | 50,000 |
|  | Cl | 66,169 |
|  | Water | 14,668,645 |
|  | **Total particles** | **14,784,814** |

Movie S1.

A 200 ns trajectory of the all-atom simulation of M dimer that includes a transmembrane domain together with a lipid bilayer and an endodomain.

Movie S2.

An example of the unstable trajectory of an envelope model containing truncated S trimer in molecular composition C2.

Movie S3.

Envelope model M1 (containing truncated S trimer) in molecular composition C1 that was run for 4μs. The trajectory provides evidence for agglomeration of M dimers into filament-like structures. Lipid molecules are depicted in sapphire blue, E pentamers in ruby red, M dimers in silver, and S trimers in gold. The model turns counter-clockwise for the first half of the movie and clockwise for the second half of the movie.

Movie S4.

Comparing the outer and inner surfaces of the envelope model M1 (containing truncated S trimer) in molecular composition C1 during a 4μs simulation run. In contrast to the outer surface of the envelope where TM domains of M dimers can be viewed forming filament-like structures, the inner surface of the envelope shows tight packing of ED domains of M dimers. Lipid molecules are depicted in sapphire blue, E pentamers in ruby red, M dimers in silver, and S trimers in gold.

Movie S5.

Envelope model M2 (containing full S trimer) in molecular composition C1 that was run for the first 1μs. The agglomeration of M dimers into filament-like structures is still apparent during this, shorter, run. Lipid molecules are depicted in sapphire blue, E pentamers in ruby red, M dimers in silver, and S trimers in gold. The model turns counter-clockwise for the first half of the movie and clockwise for the second half of the movie.

Movie S6.

A visualization of 13 μs coarse-grained molecular dynamics trajectory of a flat bilayer system that includes 41 M dimers with randomly assigned initial positions in the bilayer. Lipids are colored red/magenta; proteins are colored green; carbohydrates are colored purple.

Movie S7.

Network dynamics of domain-domain interaction network between the transmembrane (TM) domains of M dimers during 4μs simulation run. The representation of the envelope model is converted from 3D to 2D using Mercator projection, that preserves local directions and shapes. Each node represents a TM dimer, and each edge in the network represents a physical interaction between TM domains of a pair of M dimers. Newly formed edges are highlighted in orange.

Movie S8.

Network dynamics of domain-domain interaction network between the endodomains (ED) of M dimers during 4μs simulation run. The representation of the envelope model is converted from 3D to 2D using Mercator projection, that preserves local directions and shapes. Each node represents an ED dimer, and each edge in the network represents a physical interaction between ED domains of a pair of M dimers. Newly formed edges are highlighted in orange.
